# Accurate prediction of thermoresponsive phase behavior of disordered proteins

**DOI:** 10.1101/2025.03.04.641540

**Authors:** Ananya Chakravarti, Jerelle A. Joseph

## Abstract

Protein responses to environmental stress, particularly temperature fluctuations, have long been a subject of investigation, with a focus on how proteins maintain homeostasis and exhibit thermoresponsive properties. While UCST-type (upper critical solution temperature) phase behavior has been studied extensively and can now be predicted reliably using computational models, LCST-type (lower critical solution temperature) phase transitions remain less explored, with a lack of computational models capable of accurate prediction. This gap limits our ability to probe fully how proteins undergo phase transitions in response to temperature changes. Here, we introduce Mpipi-T, a residue-level coarse-grained model designed to predict LCST-type phase behavior of proteins. Parametrized using both atomistic simulations and experimental data, Mpipi-T accounts for entropically driven protein phase separation that occurs upon heating. Accordingly, Mpipi-T predicts temperature-driven protein behavior quantitatively in both single- and multi-chain systems. Beyond its predictive capabilities, we demonstrate that Mpipi-T provides a framework for uncovering the molecular mechanisms underlying heat stress responses, offering new insights into how proteins sense and adapt to thermal changes in biological systems.

## Introduction

Proteins have been shown to phase separate and organize into biomolecular condensates, which are dynamic, membraneless organelles that play pivotal roles in cellular functions [1–4]. Condensates have been shown to form in response to changing environmental stimuli, such as temperature, pH, and ionic salt concentrations [5–10]. Among various environmental factors, temperature stands out as a key regulator of protein phase behavior, influencing the adaptive assembly and disassembly of proteins. For example, proteins have been shown to self-assemble in response to heat stress to maintain cellular homeostasis [11, 12]. In contrast, inability of proteins to adapt to periods of heat stress may cause the cell to become aberrant, moving to a toxic state [6, 13]. Beyond understanding the heat stress response, predicting temperature-induced protein phase separation is advantageous particularly in materials design. For example, we can exploit thermoresponsive proteins to engineer smart biomaterials, which offer straightforward control of tunable phase transitions for applications in drug delivery, tissue engineering, and stimuli responsive fibers, surfaces, and gels [14–16]. Although there are significant opportunities for understanding adaptation and materials design, our ability to predict protein phase behaviors in response to temperature remains a major challenge. Emerging evidence suggests that the propensity for proteins to phase separate in response to temperature is not random but is encoded in their amino acid sequences [17–20]. Thus, this work focuses on extending our capabilities to predict how protein sequences encode thermoresponsive phase behavior.

Proteins exhibit different types of phase separation behaviors in response to temperature. Some proteins exhibit UCST-type (upper critical solution temperature) phase behavior, forming condensates at low temperatures and mixing into solution at higher temperatures. Other proteins exhibit LCST-type (lower critical solution temperature) phase behavior, solvating at lower temperatures and condensing at higher temperatures. UCST-type phase behaviors have been well studied, with extensive research conducted to understand how protein sequences encode UCST-type phase behaviors [18, 21–23]. These studies have reached the consensus that “UCST proteins” are enriched in polar and aromatic residues and that their ability to form condensates is driven by strong intermolecular interactions between proteins [22, 24]. As a result, significant progress has been made in developing computer models that can predict UCSTs (or upper cloud point temperatures; UCPTs) of proteins with high fidelity [25–36]. In contrast, our ability to understand and predict LCST-type phase behavior of proteins has lagged behind. This is predominantly due to the challenges in navigating complexities of solvent effects that drive LCST-type phase transitions. However, with increasing investigation into the mechanics of biomolecular condensates, there has been a new surge of interest in elucidating LCST-type phase transitions. Here, we aim to understand how protein sequences encode LCSTs and to exploit these heuristics for modeling and predictive capabilities.

Key experimental works have explored connections between protein sequence and LCST-type phase behaviors. Seminal work by Quiroz and Chilkoti investigated phase behavior of proteins identified as LCST or UCST [18]. Their study found that there are key sequence factors that may predispose a protein to exhibit an LCST or UCST. Additionally, their work demonstrated that “LCST proteins” are enriched significantly in hydrophobic residues and that hydrophobicity is a key driving factor in LCST-type phase transitions. However, using solely experimental techniques to uncover the sequence space of thermoresponsive proteins can be labor intensive and expensive. Furthermore, while experimental approaches provide valuable data, they often lack the resolution necessary to capture sub-molecule interactions and dynamics that govern protein behavior. As a result, these approaches are limited inherently in observing transient conformations, intra- and intermolecular interactions, or the precise role of solvent effects, which are critical for understanding temperature-induced phase transitions. These limitations underscore the need for computational approaches that can offer detailed insights into the mechanisms driving such phenomena.

To address this need, molecular simulations and computational modeling have proven to be advantageous, particularly in enabling the identification and exploration of the chief mechanisms at play in protein phase separation. However, capturing LCST-type phase behavior in simulations is very difficult as most models have been developed to describe UCST-type phase transitions. Atomistic simulations have provided detailed insights into single-chain properties, such as changes in radius of gyration, under varying thermal conditions [37–39], and are effective particularly in studying shorter protein chains. For instance, replica exchange Monte Carlo simulations have been employed to parameterize models to account for temperature dependence through solvation free energy values [39, 40]. However, such approaches are computationally intensive, limiting their scalability to larger systems. When studying phase separation, which requires simulating interactions among many proteins over larger length scales, atomistic simulations become increasingly computationally expensive. To overcome these limitations, residue-level coarsegrained simulations have proven to be invaluable tools. Coarse-grained approaches attempt to strike a balance between molecular resolution and computational efficiency, enabling the simulation of protein behavior over extended length scales. For example, in previous work, Dignon et al. developed a residue-resolution model for identifying protein sequences with LCSTs [41]. Building on the accuracy of all-atom simulations and the efficiency of coarse-grained models, we aim to develop a model that captures single- and multi-chain LCST-type phase behavior with both precision and efficiency.

Here, we present Mpipi-T: a physics-based residue-level model designed to capture LCST-type phase behavior of proteins accurately. Disordered protein regions have been shown to play a pivotal role in condensate formation and have been engineered for thermoresponsive properties [39, 42–44]. Therefore, Mpipi-T is designed specifically to capture the behavior of disordered proteins, addressing key challenges in accurately modeling their thermoresponsive phase transitions. Furthermore, a critical aspect of such systems is that they must achieve a certain hydration threshold in order to exhibit LCST-type properties [45, 46]. Accordingly, Mpipi-T is designed to simulate proteins in aqueous solution for the temperature range of liquid water (273 K to 373 K); ensuring that our simulations capture effects due to hydration that are essential for LCST-type phenomena. Our approach builds upon the Mpipi model, which has demonstrated good predictions for UCST-type phase behavior of disordered proteins [25]. Mpipi-T extends these capabilities to LCST-type phase behavior. By training model parameters against atomistic simulations and experimental data, Mpipi-T accurately reproduces temperature-dependent single chain and collective behaviors of disordered proteins. Leveraging Mpipi-T, we demonstrate that three key proteins—in mammals, yeast, and plants—implicated in heat stress responses all encode LCST-type phase transitions, strongly suggesting that the heat stress response is inherently LCST-type. Collectively, Mpipi-T represents a potentially valuable approach for predicting LCST-type phase transitions in proteins, elucidating underlying mechanisms of heat stress responses, and engineering thermoresponsive synthetic proteins.

## Results

### Designing a coarse-grained model for capturing single chain and collective LCST-type phase behavior

The parent model of Mpipi-T is the multiscale π–π, or Mpipi model [25]. Mpipi is a residue-level coarse-grained model that groups each amino acid into a coarse-grained bead, assigning each bead a charge, mass, and interaction parameters. Potential energy is composed of bonded, electrostatic, and short-range nonbonded pairwise interaction terms. In particular, Mpipi accounts for the dominance of π–π and hybrid cation–π interactions in driving UCST-type phase behavior of disordered proteins [47]. In doing so, Mpipi has achieved quantitative agreement with experiments for single-molecule properties near 300 K, as well as for predictions of UCST-type phase behavior of proteins. However, Mpipi does not account explicitly for temperature-dependent changes in protein solvation, which are crucial for capturing LCST-type phase behavior.

Here, our goal is to design a model that captures LCST-type phase behavior of proteins. Whereas UCST-type phase behavior is an associative phase transition driven by short-range attractions between protein residues, the LCST-type phase transition is more complex as it arises from segregative phase separation driven by entropic incompatibility between solvent and protein residues. To model LCST-type phase transitions in an implicit solvent model, it is essential to account for both temperature-dependent electrostatic interactions, which occurs due to variations in ionic screening ability, and the aforementioned temperature-driven segregative phase separation. Mpipi-T aims to capture the temperature-dependent electrostatic effects by using an analytical approach to account for changes in long-range electrostatics with temperature and a combined top-down–bottom-up approach to tune pairwise short-range nonbonded interactions as a function of temperature.

Following Mpipi, Mpipi-T is also a residue-level coarsegrained model—grouping each amino acid into a coarse-grained bead with an assigned charge, mass, and interaction parameters (Fig. 1). Potential energy encompasses bonded, electrostatic, and pairwise short-range nonbonded interaction terms. Bonded interactions are modeled using a harmonic potential, electrostatics are modeled using the Yukawa potential, and short-range nonbonded interactions are modeled using the Wang–Frenkel potential [48]. However, in deviation from the Mpipi model, the electrostatic screening interactions are now scaled as a function of temperature, and the crux of the Mpipi-T model is that we scale interaction energies in the Wang–Frenkel potential as a function of temperature based on functional forms which we design and optimize.

**FIG. 1:**
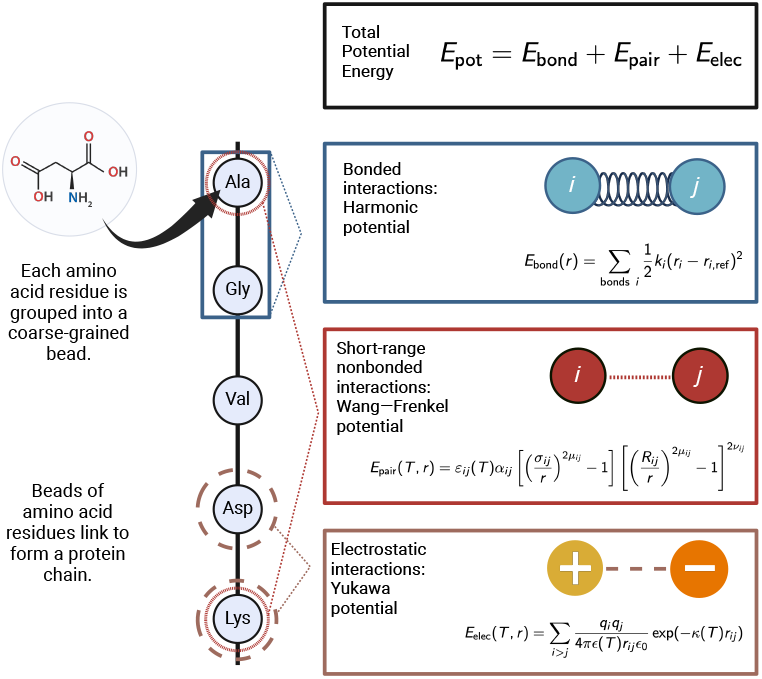
Mpipi-T is developed to capture LCST-type phase behavior by incorporating temperature-dependent functionals into the force fields equations. Pairwise *ε*_*ij*_ values (well depth) in the Wang–Frenkel potential are scaled to incorporate temperature dependence using atomistic simulations and experimental data. In the Yukawa potential, the *κ* (inverse Debye length) and *ϵ* (dielectric constant) terms are scaled analytically as a function of temperature. Figure created using BioRender.

### Deriving a temperature-dependent electrostatic potential

To account for the change in electrostatic interactions as a function of temperature, we scale analytically the *ϵ* (dielectric constant) and *κ* (inverse Debye length) terms in the Yukawa potential. To tune the dielectric constant, we use the relation:

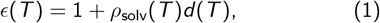

where *ϵ* is the dielectric constant, *d* is a solvent dependent parameter, and *ρ*_solv_ is the density of the solvent [49]. The solvent dependent parameter is determined as a function of temperature using experimental observations; for the temperature range of liquid water (273 to 373 K), the linear relationship of *d* and T suffices [49, 50]. The density of the solvent (water) is determined using the Kell formulation for density of water as a function of temperature [51].

The equation for the square of the inverse Debye length is:

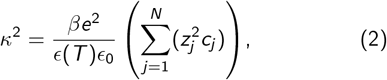

where *z*_*j*_ is the valence of ions, *ϵ* is the permittivity of the solvent, *ϵ*_0_ is the vacuum permittivity (8.85 × 10^*−*12^ C^2^ N^*−*1^ m^*−*2^), *e* is the fundamental charge (1.6 × 10^*−*19^ C), and *c*_*j*_ is the ion concentration. We scale *ϵ*(*T*) based on the description above and account for temperature in *β*, which is 1*/k*_B*T*_.

### Rationale for reparametrization of Wang–Frenkel interaction energies

While LCST-type phase behavior has been understudied relatively, previous work has demonstrated that proteins exhibiting LCSTs are enriched significantly in hydrophobic residues and that hydrophobicity is a key driving factor in the LCST-type response [18]. More specifically, as temperature increases, it becomes increasingly unfavorable for water molecules to organize around hydrophobic, nonpolar residues. Thus, the entropic penalty of water molecules being bound around hydrophobic residues drives the system to phase separate spontaneously in an endothermic process, thereby enabling escape from being trapped in an unfavorable hydration shell [16, 52, 53]. We hypothesize that as temperature is increased, the difference in magnitude of translational entropy between water molecules and proteins could result in water molecules being trapped amidst protein chains. This, coupled with disruption of van der Waals enthalpic interactions due to an increase in orientational entropy of molecules, may drive demixing of water and proteins, resulting in protein condensates in an LCST-type response.

By parametrizing the ABSINTH model [54] with reference to free energy of solvation data, Zeng et al created a robust temperature-dependent atomistic model (ABSINTH-T), which they leverage to design LCST sequences [39]. To guide LCST sequence design, they employed a genetic algorithm to generate 241 distinct sequences based on the xPxxG motif (where P is proline, G is glycine, and X is a guest amino acid residue), a pattern found commonly in ELPs and used frequently in engineering LCST proteins. By analyzing the resulting sequences, they determined the probability of different residues occupying the 1st, 3rd, and 4th positions. Notably, only five hydrophobic residues— alanine, valine, isoleucine, leucine, and methionine—had probabilities exceeding 0.1 in any of these positions, reinforcing the established link between LCST-type phase behavior and hydrophobic residue enrichment. This result aligns with analyses from Quiroz and Chilkoti, where a marked increase in enrichment of hydrophobic residues for LCST sequences compared to UCST sequences was reported [18, 23].

Based on the aforementioned insights on sequence drivers, we incorporated temperature dependence into the interaction energies for residue pairs which contained at least one of the hydrophobic residues (see Methods and Supporting Information). The key rationale behind this choice is that for other types of residues, we hypothesize that Mpipi is already able to capture temperature-dependent effects, such as the weakening of associative interactions due to accelerated kinetics. However, for hydrophobic residues, the implicit solvent aspect of Mpipi is not able to account for the entropic penalty of caged water molecules which drives LCST-type phase transitions. Furthermore, temperature-dependent behavior of charged residues is already accounted for in the electrostatic potential.

### Revised function for Wang–Frenkel potential

When rescaling the original Mpipi *ε*_*ij*_ values, some of the new *εij* values become negative when scaled down as temperature decreases. The original Wang–Frenkel potential equation [48] models interactions based on an attractive well-depth *ε*_*ij*_ (Eq. 3):

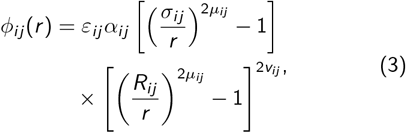

where

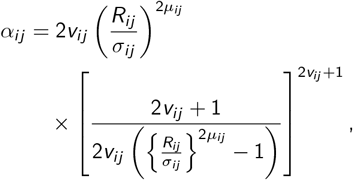

*ϕ*_*ij*_ (*r*) is the potential function, *ε*_*ij*_ is the well depth, *σ*_*ij*_ is the distance where the intermolecular potential is 0, and *R*_*ij*_ = 3*σ*_*ij*_ is the distance where *ϕ*_*ij*_ (*r*) vanishes. *µ*_*ij*_, and *ν*_*ij*_ are additional parameters used to control the shape and steepness of the potential.

However, when *ε*_*ij*_ for a pair of beads becomes negative, the interaction between the beads becomes increasingly attractive (approaching ∞) as they move closer together, which would be unphysical (Fig. 2). To address this discrepancy, we create two versions of a revised potential that expresses repulsive interactions for bead pairs with negative *ε*_*ij*_ (Eqs. 4 and 5):

**FIG. 2:**
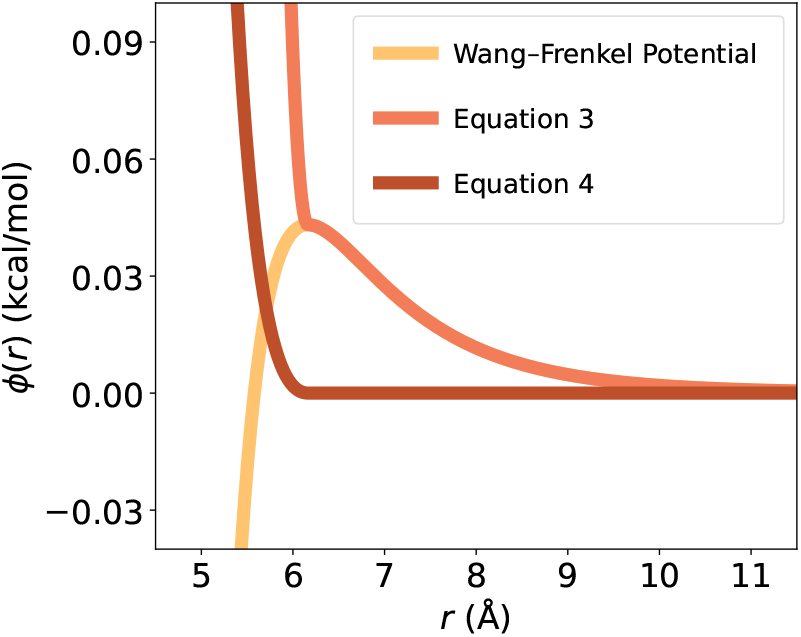
New implementations of the Wang–Frenkel potential account for purely repulsive interactions. Based on the functionals, Equation 4, similar to the WCA potential, preserves the bead size for an amino acid; hence is chosen in the implementation for the Mpipi-T model.

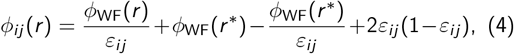

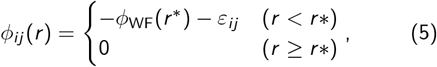

where 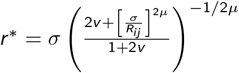 and *ϕ* _WF_ (*r*) is the canonical Wang–Frenkel potential.

Eq. 4 is similar to the Ashbaugh–Hatch implementation of the Lennard-Jones potential [55], and Eq. 5 is similar to the WCA potential [56]. We choose Eq. 5, since this implementation preserves the molecular diameter (i.e., the bead size). The modified Wang–Frenkel potential has been implemented in the LAMMPS software [57], which is used for carrying out simulations with Mpipi-T (see Methods).

### Creating an initial set of functional forms for temperature-dependent Wang–Frenkel interaction energies

To generate the functional forms for *ε*_*ij*_, we create a fitness function to optimize the model:

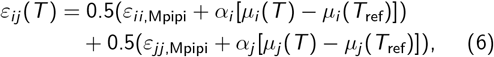

where *i* and *j* are the two amino acids, *µ* is a temperature-dependent function with variable parameters, *T*_ref_ = 298.15 K is the reference temperature, *α* is a variable parameter, *ε*_*XX*,Mpipi_ is the well depth energy from the original Mpipi model. *µ*_*X*_ is set as a parabolic function *µ*_*X*_ (*T*) = *a*_*X*_ *T* ^2^ + *b*_*X*_ *T* + *c*_*X*_, where *a, b, c* are variable parameters and *T* is temperature in Kelvin.

We set the initial parameters *a, b, c* in *µ*_*X*_ based on data from atomistic simulations using AMOEBA (polarizable force field) [60] of free energy of solvation as a function of temperature for amino acid side chain moieties [39].

### Tuning functional forms using a top-down approach

To tune the functional form parameters for all of the pairs that included a hydrophobic residue, we use experimental data of cloud point temperatures for LCST protein sequences [18]. Since the experimental data is obtained from the dilute regime and computer simulations of phase coexistence are more precise in the dense regime (i.e., less statistical uncertainty), we devise a new method CLOUD-FIT for using slab simulations to calculate cloud point temperatures of proteins (see Methods and SI for details). In short, our approach simulates slabs at the experimental concentration and exploits finite-size effects to identify density changes as a function of temperature and map out the “density versus temperature curve” for the dilute regime, thereby extracting the cloud point temperature (Fig. 3a).

**FIG. 3:**
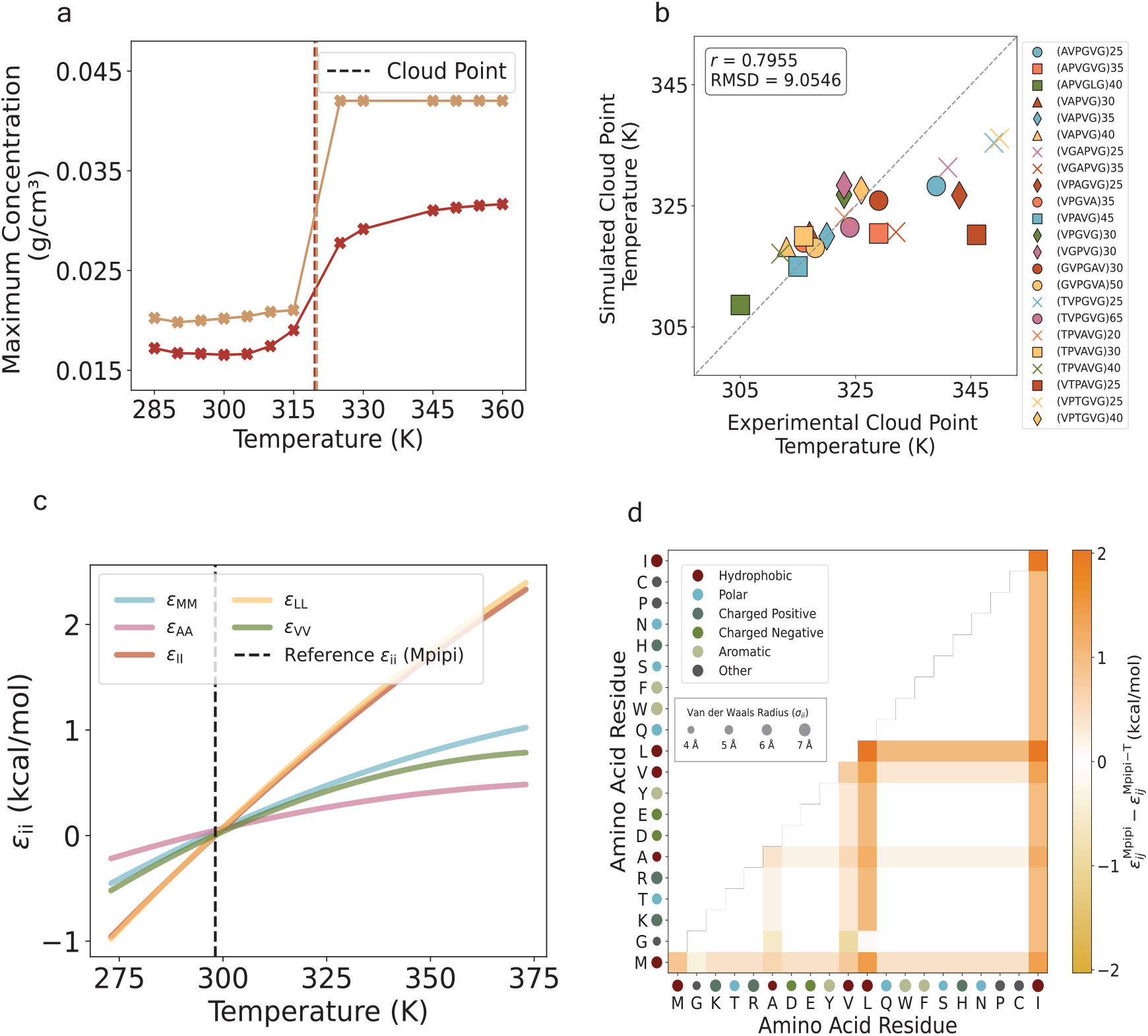
By optimizing with atomistic simulations and experimental data, Mpipi-T is trained to capture LCST-type phase behavior. (a) By devising and implementing a new method CLOUD-FIT, the cloud point is found from concentration profiles of proteins simulated in the dilute phase (two examples are shown here). (b) The optimized model is chosen based on the lowest RMSD value of the simulated and experimental cloud point temperatures. The dataset is divided into three blocks, and each data point represents the mean of the blocks. The error bars are computed using the standard error, but they are not shown as they are smaller than the size of the data points on the graph. (c) A subset of functional forms for homotypic interactions between hydrophobic residue pairs as well as (d) the contact map for interaction energy differences between Mpipi and Mpipi-T at 360 K is shown.

After the optimization, we converge on three models which performed the best. We choose the model that had the lowest RMSD value between the simulated and experimental cloud point temperatures (Fig. 3b). In addition to this optimized model, we develop two other models (Figs. S1a, S1b) that also capture the experimental data well. We recommend benchmarking the systems with all 3 models, and then moving forward with the one that is best suited for the test system. Our final set of parameters that produce the functional forms for each *ε*_*ij*_ in our optimized model (Fig. 3c, 3d) for *ε*_*ij*_ are given in Tables S2–S21 in the SI.

### Testing ability of the optimized model to distinguish between LCST- and UCST-type phase behavior

After optimizing Mpipi-T, we evaluate whether our model is able to distinguish between LCST and UCST proteins. A cheap way to test this is by computing the Flory scaling exponent over a range of temperatures (see Methods) for a set of LCST and UCST sequences from experiments [18]. As expected, Mpipi is able to predict UCST-type but not LCST-type phase behavior (Figs. 4a, 4c). In contrast, Mpipi-T is able to distinguish UCST sequences from LCST ones (Figs. 4b, 4d). Mpipi-T, therefore, preserves the ability to predict UCST-type phase behavior, while capturing LCST-type phase transitions.

**FIG. 4:**
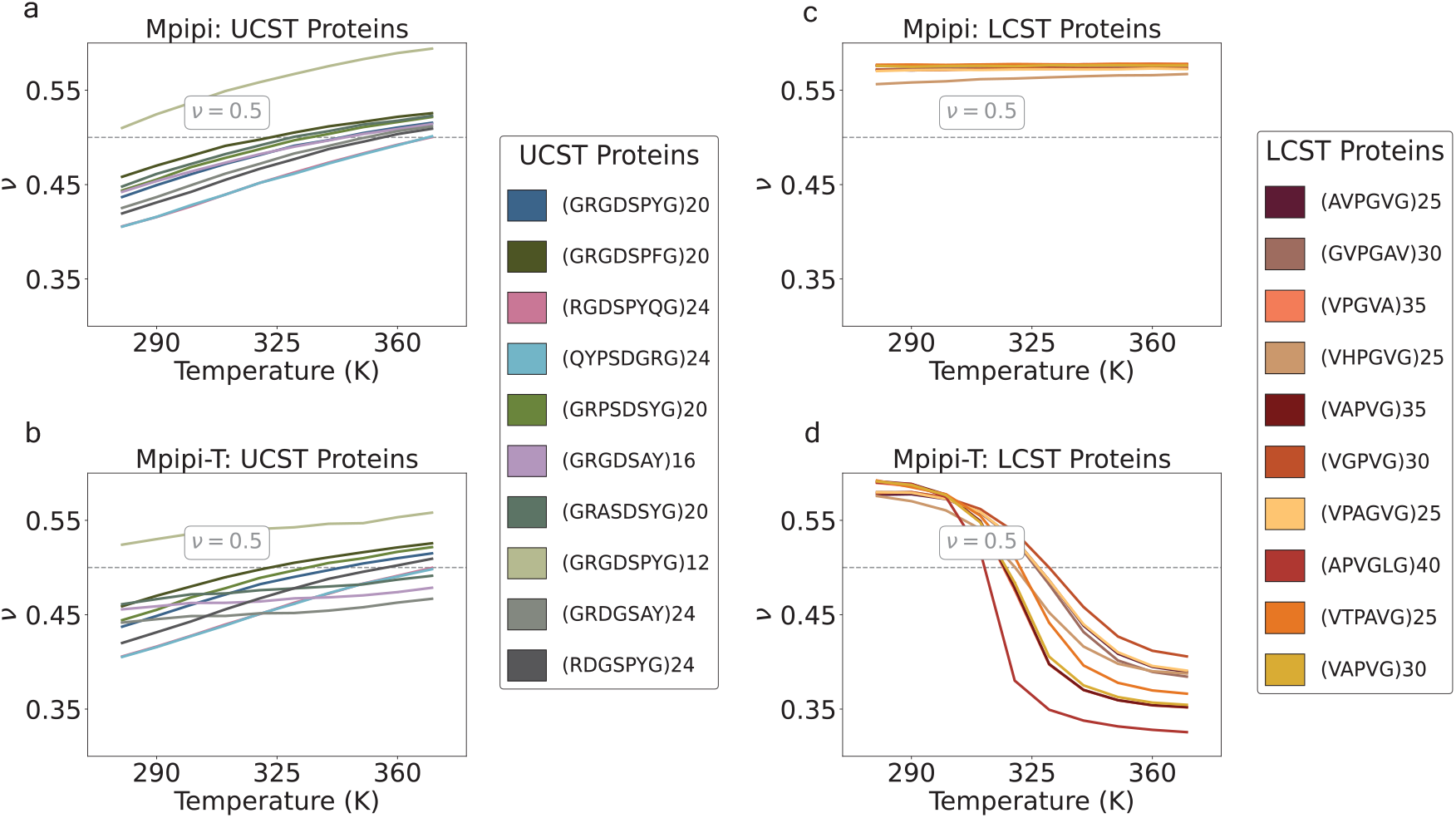
Mpipi-T captures LCST-type phase behaviors, while retaining the ability to predict UCST-type phase behavior as well. The Flory exponent (*ν*) is computed to test whether the model is able to distinguish between UCST: a) Mpipi, b) Mpipi-T, and LCST: c) Mpipi, d) Mpipi-T sequences. For each set of proteins, the UCST and LCST protein sequences are shown in the legends on the right-hand panels. While both Mpipi and Mpipi-T are able to predict UCST transitions, only Mpipi-T is able to predict LCST transitions. To evaluate the error, the dataset is divided into three blocks, with the mean of the blocks used to determine the data points. The error bars are computed using the standard error, but they are not shown as they are smaller than the width of the lines. The dashed gray line indicates *ν* = 0.5, which represents the coil-to-globule transition temperature.

### Further validation of optimized model against ELP sequences

Finally, we characterize the phase behavior of a set of 5 ELP sequences—for which we have experimental data of the dilute phase density as a function of temperature (left arm of the binodal)—to evaluate the performance of Mpipi-T [61]. Since the sequences are quite long (ranging up to 800 residues in length), we initially rely on the single chain dynamics to provide insight into whether Mpipi-T predicts a strong agreement between the simulated coil-to-globule transition temperature (*T*_θ_) and the estimated experimental critical temperature (*T*_c_). The full phase diagrams and single-chain correlations for all models are available in the SI (Figs. S2–S5). In comparison to the HPS-T model [41], which we observe overpredicts transition to the globule state (Fig. 5a), Mpipi-T predicts the correct range for the coil-to-globule transition as well as achieves a high Pearson correlation with estimated experimental critical temperature (Fig. 5b).

**FIG. 5:**
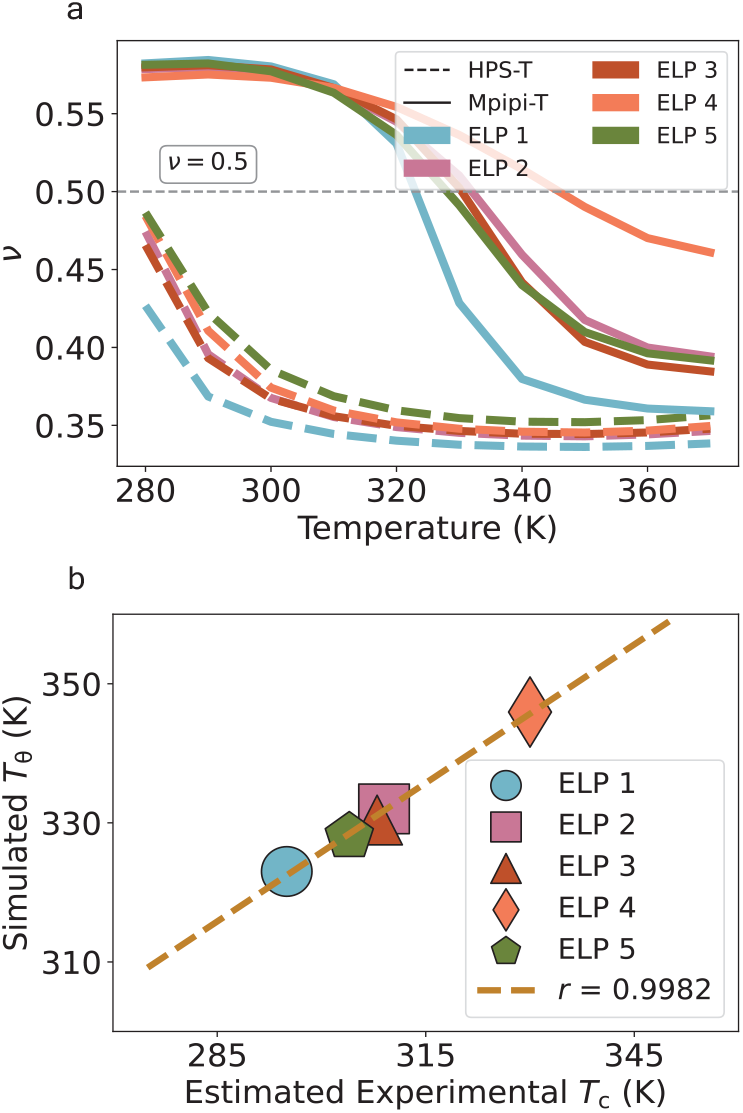
Mpipi-T achieves good predictions for an experimental dataset of protein sequences. (a) Flory scaling exponent versus temperature for ELP sequences using HPS-T and Mpipi-T models. The dashed gray line in indicates *ν* = 0.5, which represents the coil-to-globule transition temperature. To evaluate the error, the dataset is divided into three blocks, with the mean of the blocks used to determine the data points. The error bars are represented by the standard error, however they are not shown as they are smaller than the width of the lines. b) Simulated coil-to-globule transition temperature (*T*_θ_) versus simulated experimental critical temperature for a set of ELP proteins. The brown line represents the line of best fit, and the Pearson correlation is shown in the inset. Compared with HPS-T, which overpredicts the LCSTs, (b) Mpipi-T achieves a high Pearson correlation value for comparing coil-to-globule transitions with estimated experimental critical temperatures. The dataset is divided into three blocks, with the mean of the blocks used to determine the data points. The error bars, representing the standard error, are not shown as they are smaller than the size of the data points.

### Generating phase diagrams for proteins implicated in heat stress responses

In addition to validation of Mpipi-T, we test whether Mpipi-T predicts LCST-type phase transitions for proteins that have been shown experimentally to form condensates under heat stress. We focus on proteins found in three different organisms, exploring the predictive capabilities of our model in diverse biological contexts (Fig. 6).

**FIG. 6:**
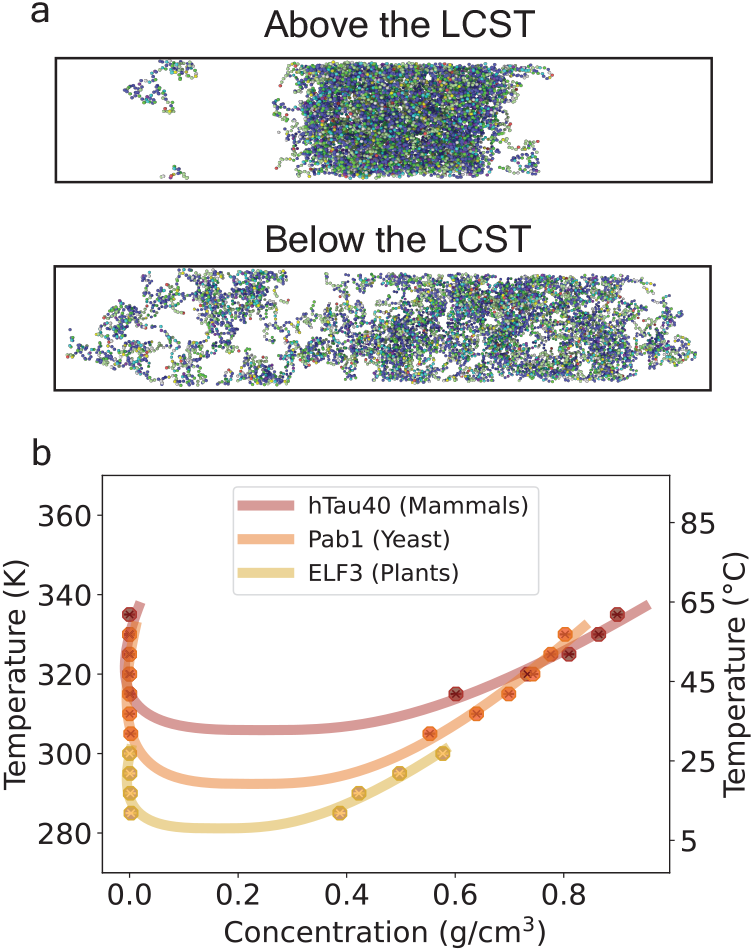
Mpipi-T predicts LCST transitions for proteins involved in heat shock responses. Three proteins that condense under heat stress are tested: hTau40 (Alzheimer’s disease associated in humans), Pab1 (stress granule marker in Baker’s yeast), and ELF3 (circadian rhythm and flowering regulator in mouse-ear cress). The respective sequences simulated were the 2N4R isoform of human tau (K18 variant, or residues 244 to 372) [58], the low-complexity P domain of Pab1 (residues 419 to 502) [7], and the prion-like domain of ELF3 (residues 430 to 609) [59]. (a) A visual for proteins phase separating above the LCST is shown in the top panel and proteins mixing below the LCST is shown in the bottom panel. (b) Phase diagrams for each of the proteins are created based on the simulation data. The data points are obtained from direct coexistence simulations at various temperatures to determine dense and dilute concentrations. These data are then fitted using the law of coexistence densities and the law of rectilinear diameters to construct the phase diagrams and identify critical temperatures. Each trajectory is divided into three blocks, with the mean of the blocks used to determine each data point. The error bars, representing the standard error, are not shown as they are smaller than the size of the data points.

#### Mammalian hTau40

For mammals, we test hTau40, the longest isoform in the central nervous system of the Alzheimer’s disease-associated Tau protein in humans. The hTau40 protein expresses pathological abnormalities in the entorhinal cortex and synaptic circuits during the early stages of Alzheimer’s disease [62]. Understanding how temperature sensitivity and mutations influence hTau40 phase behavior could provide insights into the correlation between these factors and disease onset or progression. Interestingly, experimental work has demonstrated LCST-type phase behavior and phase separation of hTau40 under heat stress [5, 58, 63, 64]. By leveraging simulations, we bridge the gap left by experimental work, extending these studies by computing the temperature-dependent phase diagram of hTau40. Using Mpipi-T, we predict an LCST for hTau40 at 305.9 K. This value aligns closely with human body temperature of 310.15 K, reinforcing the biological relevance of our predictions. However, this LCST specifically corresponds to simulations conducted on the K18 region, which consists of the four imperfect repeats that are known to drive phase separation. Given their role, it is reasonable that the LCST for K18 is lower than physiological temperature. This suggests that these repeats alone undergo phase separation more readily. However, this does not imply that full-length hTau40 is always phase-separated under physiological conditions. In fact, additional simulations which we performed (not shown) of full-length hTau40 suggest a slightly higher LCST of 309.5 K, indicating that the entire protein exhibits a marginally greater resistance to phase separation than K18 alone and that full length hTau40 may be near criticality in living cells.

#### Yeast Pab1

For yeast, we model Pab1, a core stress granule poly(A)-binding protein in Saccharomyces cerevisiae. The phase behavior of Pab1 under heat stress has been studied extensively, and this protein serves as a valuable marker for understanding stress granules in a eukaryotic model organism [6, 7, 65]. Characterizing the phase boundaries of Pab1 is important for understanding thermoresponsive fitness in organisms and sequence-encoded evolutionary adaptations to environmental conditions. Experimental studies have demonstrated that disordered Pab1 undergoes phase separation under heat stress and have predicted an LCST using pH–temperature phase diagrams [7, 42]. Here, we use simulations to generate a phase diagram along the temperature–density plane and predict that Pab1 does in fact have an LCST at 292.4 K. This value corresponds closely to the lowest demixing temperature observed in a pH and temperature phase diagram of purified Pab1 [7], adding confidence in our predictions.

#### Plants ELF-3

For plants, we examine ELF-3 (Early Flowering 3), a large scaffold protein that is part of the evening complex in Arabidopsis thaliana, a model organism for plants. ELF-3 acts as a thermosensing protein; its prion-like domain unfolds in response to a decrease in temperature, triggering the activity of the evening complex [59]. This behavior is crucial for plant development, as temperature influences phenological and circadian events directly. Understanding the phase behavior of ELF-3 can provide insights into how plants respond to temperature changes, which is relevant particularly in the context of climate change. Additionally, studying homologous proteins in varying environmental conditions can offer valuable information for applications in invasive species control, crop development, and assessing the fitness of native and wild species under changing climates. Experimental work has not yet provided a full phase diagram for ELF-3, but our model predicts an LCST at 281.2 K. This prediction aligns with the temperature at which speckles have been observed to form in experimental studies [66], further validating the relevance of our computational approach.

Together, our predictions using Mpipi-T provide insights into the phase behavior of proteins found in different organisms, offering data that complement and extend experimental findings. These simulations open new avenues for understanding the environmental and evolutionary dynamics of protein phase separation, with implications for human health, stress responses in eukaryotes, and plant adaptation.

## Discussion

In this work, we demonstrate that by parametrizing temperature-dependent electrostatic interactions analytically and short-range non-bonded interactions with a combined top-down–bottom-up approach, Mpipi-T captures accurately LCST-type phase behavior of disordered proteins. Mpipi-T is built upon the Mpipi model—extending its predictive capabilities from USCT sequences to include LCST.

To develop Mpipi-T, we introduce temperature dependence in dielectric interactions by scaling analytically both the inverse Debye length and dielectric constant as functions of temperature. Additionally, we implement a modified Wang–Frenkel potential to incorporate repulsive interactions while preserving particle interaction sizes. A key advancement is training the interaction parameters, specifically the well depth energy, in the Wang–Frenkel potential to capture temperature effects using both atomistic simulations [39] and experimental data [18] for each amino acid pair. Upon testing, Mpipi-T retains the UCST predictive capabilities of its parent model, while extending its scope to distinguish proteins exhibiting LCST-type phase behavior. Finally, when applied to ELP sequences with LCST-type phase transitions, previously characterized experimentally, Mpipi-T demonstrates strong agreement with experimental observations, reinforcing its reliability in capturing LCST-type phase behavior.

Despite its strengths, Mpipi-T has certain limitations. The model does not capture protein conformational changes with temperature explicitly, which can influence phase behavior, in structured proteins particularly. While this is less critical for disordered sequences—the primary focus of this study— future extensions could incorporate chain rigidity and unfolding effects to enhance predictive power. Additionally, the scarcity of experimental data, especially for native LCST proteins, poses challenges for model training and validation. Most available data come from ELPs and synthetic sequences, highlighting the need for broader experimental benchmarks.

Notwithstanding these, Mpipi-T demonstrates potential as an approach for investigating LCST-type phase behaviors in proteins, offering molecular-level insights into their phase transitions. Mpipi-T can be used widely to probe whether proteins exhibit LCST-type phase behavior and analyze quantitatively their transition temperatures in both single- and multi-chain systems. To facilitate broader adoption, we have made all necessary input files available and implemented Mpipi-T within the open-source LAMMPS software [57] (see Code and Data Availability statement). Furthermore, Mpipi-T holds promising utility for facilitating the rational design and discovery of novel protein sequences tailored for specific thermal and environmental conditions. Mpipi-T can be harnessed to aid in the design thermoresponsive proteins that exploit biological adaptive capabilities for innovative biomaterials, including advanced clothing or sensing technologies.

Beyond engineering applications, Mpipi-T presents opportunities for probing the physicochemical underpinnings of temperature-induced phase transitions, particularly in the context of heat shock responses in living systems. Many proteins have been observed to phase separate in response to temperature changes, but conflicting views exist regarding the exact mechanisms involved in such responses. The mechanism of heat-induced condensation has long been thought to be proteotoxic stress leading to misfolding and aggregation, with the heat shock response and recovery mediated by molecular chaperones [6, 7]. However, recent studies suggest that certain proteins encode adaptive LCST-like phase transitions within their sequences, allowing them to sense directly and respond reversibly to temperature changes without the occurrence of protein damage [6, 7, 65, 67]. Here, we leverage Mpipi-T to probe three well-studied proteins implicated in heat stress responses (in mammals, yeast, and plants). Our simulations predict LCST-type phase transitions for all sequences— notably, occurring within the physiological temperature range of the respective organisms. These findings suggest that LCST-type phase behavior may be an intrinsic component of the stress response, expanding the current understanding of how cells maintain homeostasis under thermal stress. Moving forward, a broader investigation into stress-related proteins could reveal whether LCST-driven phase transitions are a widespread adaptive strategy in cellular stress responses.

## Methods

### Experimental cloud point determination

The experimental cloud point temperatures are determined using absorbance curves. The cloud point corresponds to the temperature at which the transmittance of the solution is reduced to 0.5 [68, 69], which, according to Beer-Lambert law, corresponds to an absorbance value of 0.301. The temperatures at which the absorbance reached 0.301 are extracted from the curves and recorded as the experimental cloud point [18].

### Computational cloud point determination

#### Initial approach

*cluster formation*. An initial attempt to compute cloud point temperatures involved simulating cluster formation. Simulation boxes were constructed using the protein concentrations reported in the experimental studies. These simulations were run at various temperatures, monitoring for the formation of clusters. However, this method proved expensive computationally, as clusters required significant time to form across the temperature range.

#### CLOUD-FIT

##### a new method for computing cloud point in simulations

To address these limitations, a new method, CLOUD-FIT (Cloud Point Analysis Using Finite Size Simulations) was devised and implemented. At a fixed concentration, the cloud point is the temperature at which the system becomes turbid due to phase separation. In experiments, these measurements are often conducted at low system concentrations. For LCST sequences, the sample is heated gradually until it reaches the left arm of the binodal (i.e., becomes cloudy). Since nucleation events are rare in the dilute solutions, CLOUD-FIT exploits finite-size effects to capture density fluctuations at the target system concentration. This approach uses the slab technique to allow for more efficient determination of cloud point temperatures. The methodology is described as follows:

##### System preparation

Simulation boxes with proteins are constructed at the desired system concentration. 64 (4×4×4) replicates are used if the protein length is less than 190 residues and 27 (3×3×3) replicates are used for proteins with lengths greater than or equal to 190 residues. The simulation boxes are then compressed via *NPT* simulations to create slabs of high peptide density (0.8–1.0 g/cm^3^). Subsequently, the z-dimension (or long axis) is extended to achieve the target protein concentration (that matched the experimental setup).

##### Simulation protocol

Canonical (*NVT*) simulations, using a Langevin thermostat with a relaxation time of 5 ps, are performed with 50 ns equilibration followed by a 400 ns production run (with a timestep of 10 fs) in 5 K increments over a temperature range of 275 K to 370 K. At each temperature, the density profile along the z-axis of the slab system is calculated every 1 ns.

##### Cloud point identification

The cloud point is identified by analyzing the density profiles along the z-axis. The first temperature at which the density profile exhibited a sharp spike is determined to be the cloud point, corresponding to the onset of condensate formation (see Fig. S6). The cloud point temperatures obtained from simulations are compared with experimental measurements to assess the accuracy of CLOUD-FIT in predicting experimental cloud points.

### Direct coexistence simulations

To compute phase diagrams, direct coexistence simulations are performed following a similar procedure as in Ref. 25. Given that these systems exhibit lower critical solution temperatures (LCST), simulations are conducted at temperatures above the critical temperature.

Each system is simulated in a canonical ensemble, using a Langevin thermostat with a relaxation time of 5 ps. Each simulation included a 50 ns equilibration period followed by a 400 ns production run (with a timestep of 10 fs). At each temperature, the density profile along the z-axis of the slab system is calculated every 1 ns.

To estimate critical points from the simulated data, we use the law of coexistence densities and the law of rectilinear diameters [70], as in previous work [25].

### Computing Flory scaling exponent and coil-to-globule transition temperature

To calculate the Flory scaling exponent and identify the coil-to-globule transition, an alternate relation based on the radius of gyration is used. The standard equation that depends on interresidue distance to compute Flory exponent is valid for homopolymers of infinite length; however, for heteropolymers of finite length, the alternate relation provided a more accurate description of the transition behavior.

This relation incorporates the interpenetration function in the limit of long chains and uses the Zimm (1948) relation for the second virial coefficient and the average radius of gyration for a solution of arbitrary polydispersity [71].

From this framework, the derived equation is [72]:

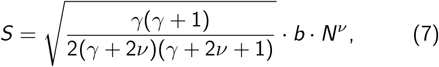

where *S* is the radius of gyration, *ν* is the Flory exponent, *b* is a prefactor set to 5.5 Å [37, 73]—resulting from the square root of the product of persistence length around 4.0 Å [74] and the distance between the C-*α* atoms set to 3.8 Å [75]—*N* is the number of bonds, and *γ* is set to 1.1615 as derived from the exponent of the Schultz distribution form for equilibrium length [76].

#### Simulation Protocol

Canonical ensemble simulations, using a Langevin thermostat with a relaxation time of 5 ps, are conducted with a 0.5 µs equilibration period followed by 3 µs of production, using a timestep of 10 fs. Simulations are performed in 10 K increments over a temperature range of 280 K to 370 K. The radius of gyration is calculated every 1000 timesteps (0.01 ns).

#### Numerical Analysis

The radius of gyration data is used in conjunction with Eq. 7 to compute the Flory scaling exponent. A numerical solver is employed to estimate the values at each temperature. The resulting data are interpolated to identify the temperature at which *ν* = 0.5, corresponding to the coil-to-globule transition [77]. Note that due to steric repulsions in an excluded volume chain, the coil-to-globule transition occurs at *ν* = 0.588 for infinite length homopolymers [78]. However, using Eq. 7, *ν* = 0.58 has been reported for random-coil protein chains that are denatured chemically [74, 79], implying that the coil-to-globule transition for finite length heteropolymers (e.g., disordered proteins) occurs at *ν <* 0.58. Thus, our data are interpolated to identify the temperature at which *ν* = 0.5, which should more closely correspond to the coil-to-globule transition.

## Code and Data Availability

The source code for running Mpipi-T in LAMMPS, input scripts, compilation instructions, and data used to produce figures and results can be found on the Joseph Group GitHub repository: https://github.com/josephresearch/Mpipi-T.

## Acknowledgments

We thank Prof. Dimitrios Fraggedakis for insightful discussions on electrostatics and polymer theory and Prof. Howard Stone for helpful feedback on the manuscript. We also thank members of the Joseph Group for their feedback during the development of Mpipi-T. J.A.J. acknowledges research support from the Chan Zuckerberg Initiative DAF (an advised fund of Silicon Valley Community Foundation; grant 2023-332391), the National Institute of General Medical Sciences of the National Institutes of Health under Award Number R35GM155259, and the National Science Foundation (NSF) through the Princeton University (PCCM) Materials Research Science and Engineering Center DMR-2011750. All simulations in this work were performed using the Princeton Research Computing resources at Princeton University, which is a consortium of groups led by the Princeton Institute for Computational Science and Engineering (PICSciE) and the Office of Information Technology.

