## Supporting Information for "Accurate prediction of thermoresponsive phase behavior of disordered proteins"

### Supporting Information for:
### Accurate prediction of thermoresponsive phase behavior of disordered proteins

Ananya Chakravarti[1,2] and Jerelle A. Joseph[1,2,*]

[1]Department of Chemical and Biological Engineering,
Princeton University, Princeton, NJ 08544, USA

[2]Omenn–Darling Bioengineering Institute, Princeton University, Princeton, NJ 08544, USA

(Dated: March 5, 2025)

#### Contents

I. Model Optimization 1

II. Model Testing 1

III. CLOUD-FIT: Cloud Point Computations 7

IV. Fitness Function Parameters for Model 3 8

References 18

#### I. Model Optimization

**Optimization of Mpipi-T Models 1, 2, and 3.** To develop Mpipi-T, we optimize three distinct models, each employing varying parametrizations of $\varepsilon$ in the Wang–Frenkel potential. To this end, the simulated cloud point temperatures (see CLOUD-FIT method in Fig. S6) are trained against experimental data for LCST protein sequences, as reported by Quiroz et al. [1]. These sequences represent a benchmark for modeling LCST phase behavior, providing a robust dataset for parameter optimization.

The cloud point optimization show that all three models achieved RMSD values between 9 K and 13 K (Fig. S1). This level of accuracy is comparable to the performance of the parent Mpipi model, which exhibited an RMSD of 9 K for the critical temperatures extracted from phase diagrams of A1-LCD wild-type and mutant proteins [2, 3]. Model 3 is chosen as the main Mpipi-T model (i.e., for results in the main text) as it has the lowest RMSD value. However, all three models should capture LCST phase behavior reliably.

#### II. Model Testing

**Testing Mpipi-T Models 1, 2, and 3 show high correlation between simulations with experiments.** To further evaluate the accuracy of the Mpipi-T models, single-chain simulations are performed for five ELP sequences using all three optimized models. These simulations are used identify the coil-to-globule transition temperatures (i.e., $T_\theta$) and compare them with the critical temperatures that are estimated experimentally (i.e., $T_c$) [4]. The sequences of the ELPs tested are as follows:

| ELP Variant | Sequence |
|---|---|
| ELP-1 | MSKGPG-$(VGPVG)_{160}$-Y |
| ELP-2 | MSKGPG-$(VPGVGVPGAG)_{40}$-Y |
| ELP-3 | $((VPGVGVPGAG(VPGVG)_4$-$VPGAG(VPGVG)_3)$-$GKG)_8$-Y |
| ELP-4 | MSKGPG-$(VPGAG)_{80}$-Y |
| ELP-5 | MSKGPG-$(VPGVG)_{40}$-Y |

TABLE S1: ELP Sequences Used for Testing

Given that some of these sequences are up to 800 residues long, direct coexistence simulations to compute phase diagrams require extensive simulation times to achieve proper equilibration. Instead, single-chain simulations are first utilized as a proxy for multi-chain simulations.

Across all three models, the simulated coil-to-globule transition temperatures exhibit a high Pearson correlation with the critical temperatures that are estimated experimentally, with correlation coefficients exceeding 0.98 (Fig. S2). Among the three models, Model 3 demonstrates the highest Pearson correlation, highlighting its superior ability in capturing the relationship between sequence and $T_\theta$ for ELP sequences.

**Computation of phase diagrams for ELP sequences to evaluate the performance of the three Mpipi-T models.** Direct coexistence simulations are used to compute phase diagrams of the five ELP sequences. The Pearson correlation coefficients be-

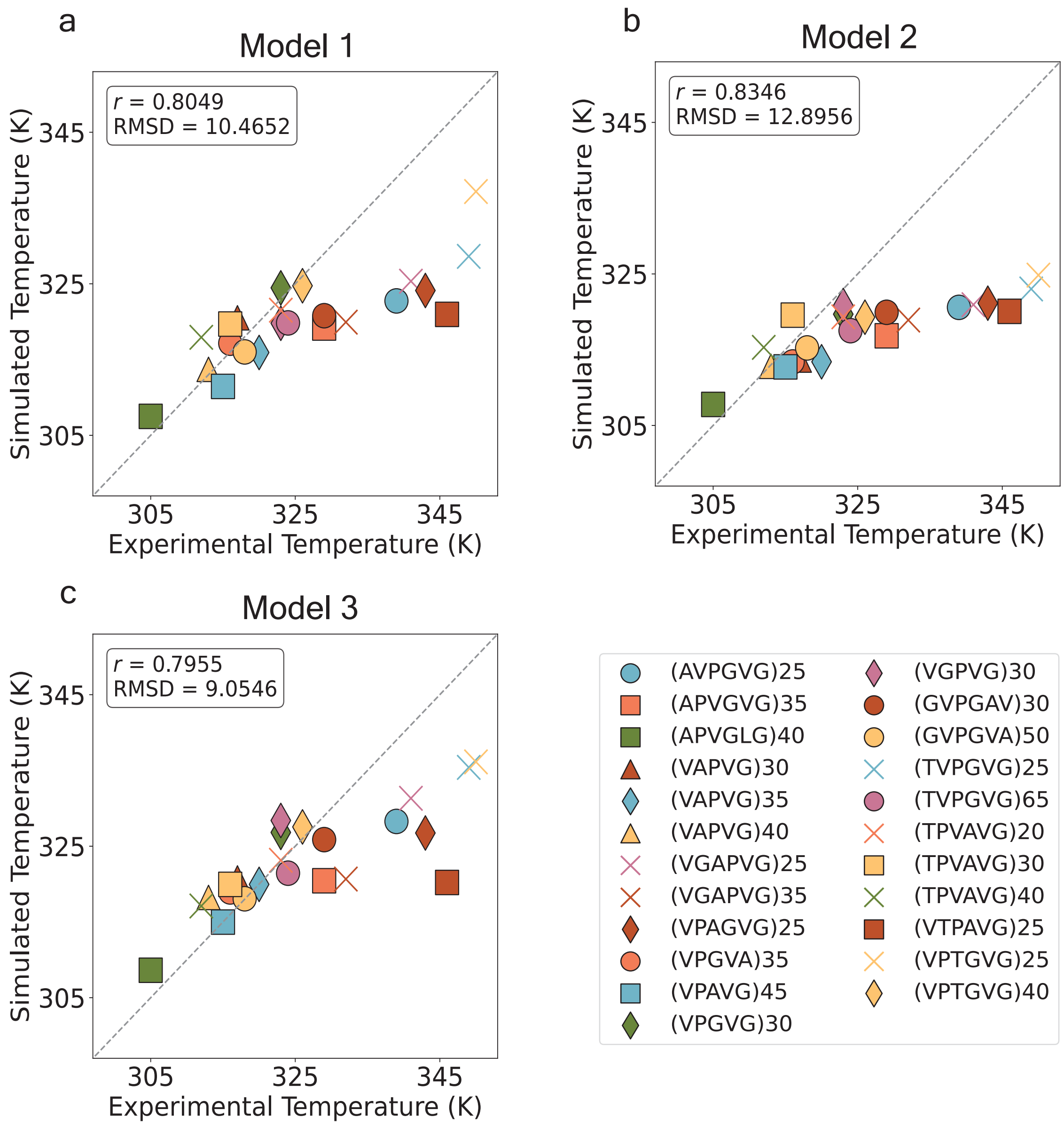


FIG. S1: **Cloud point data from optimization of Mpipi-T for** (a) Model 1, (b) Model 2, and (c) Model 3. The dataset is divided into three blocks, with the mean of the blocks used to compute the data points. The error bars, evaluated as the standard error, are not shown as they are smaller than the size of the data points. The legend, shown in the lower right panel, lists each protein sequence that was simulated. Model 3 is chosen due to its lowest value for RMSD between simulated and experimental cloud point temperatures.

tween the critical temperatures predicted by the models and the estimated values from experimental measurements are high: 0.958 for Model 1, 0.986 for Model 2, and 0.932 for Model 3. However, the root-mean-square deviation (RMSD) values reveal greater variability: 16.2 K for Model 1 (Fig. S3), 12.4 K for Model 2 (Fig. S4), and 18.9 K for Model 3 (Fig. S5). It is important to note that we estimate the experimental critical temperature by extrapolating the left arm of the binodal, as this is the available data from experiments [4].

ELP-1, which is more than 800 residues long, is a major contributor to the higher RMSD values. The large size of this sequence makes it challenging to collect reliable statistics, as achieving equilibrium in multi-chain simulations for such a system is expensive computationally.

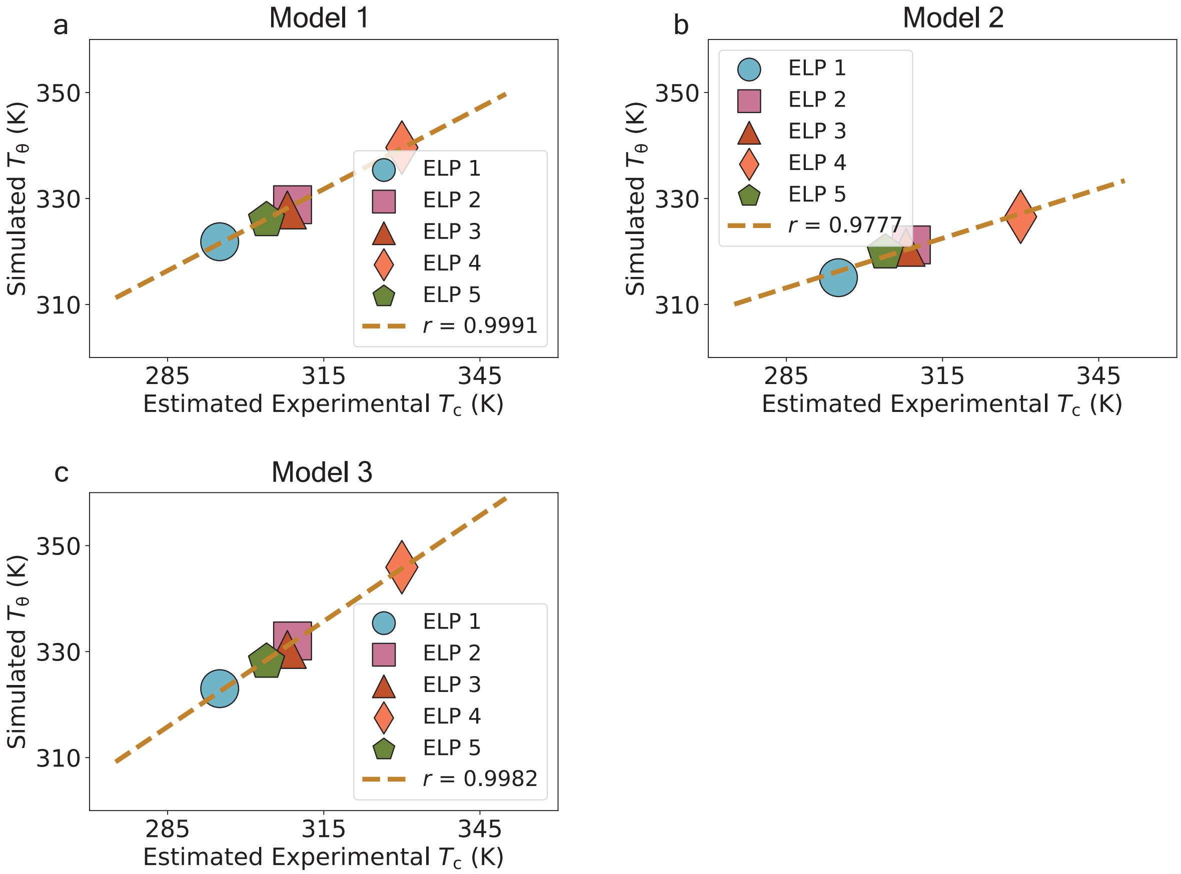


FIG. S2: **Testing of Mpipi-T models by comparing simulated single chain coil-to-globule transition to estimated critical temperature for:** (a) Model 1, (b) Model 2, and (c) Model 3. The dataset is divided into three blocks, with the mean of the blocks computed to determine the data points. The error bars, evaluated using the standard error, are not shown as they are smaller than the size of the data points. The brown line represents the line of best fit. All 3 models perform well, reflecting high Pearson correlation values.

Additionally, the critical temperature of ELP-1, estimated at approximately 295 K from experiments, is relatively low. Simulating such low temperatures requires even longer equilibration times due to slow dynamics, further complicating the reliability of the predictions.

Another factor influencing the final RMSD values is our effort to avoid overfitting. Experimental data can vary between trials; however, we only have access to the average or one of the trials. Furthermore, the experimental data used to test the models are sourced from a study different from the data used for parameter optimization. Variations in experimental methodologies may introduce discrepancies in critical temperature measurements, reflecting their inherent variability.

Despite these challenges, the high Pearson correlation values indicate that the Mpipi-T models capture the overall trends in phase behavior for the ELP sequences effectively. Here, Model 2 performs best, balancing accuracy and reliability, as evidenced by its highest Pearson correlation and lowest RMSD values.

Overall, these results demonstrate that all three Mpipi-T models are useful, but their suitability may depend on the context and the specific system being simulated. For example, Model 3 performs best in capturing LCST behavior across a broad range of disordered sequences and describing the coil-to-globule transition of long ELPs. However, Model 2 shows slightly better performance in reproducing experimental phase diagrams of long ELPs (test here). In summary, we encourage users to test all three models and select the Mpipi-T model that best suits their simulation needs.

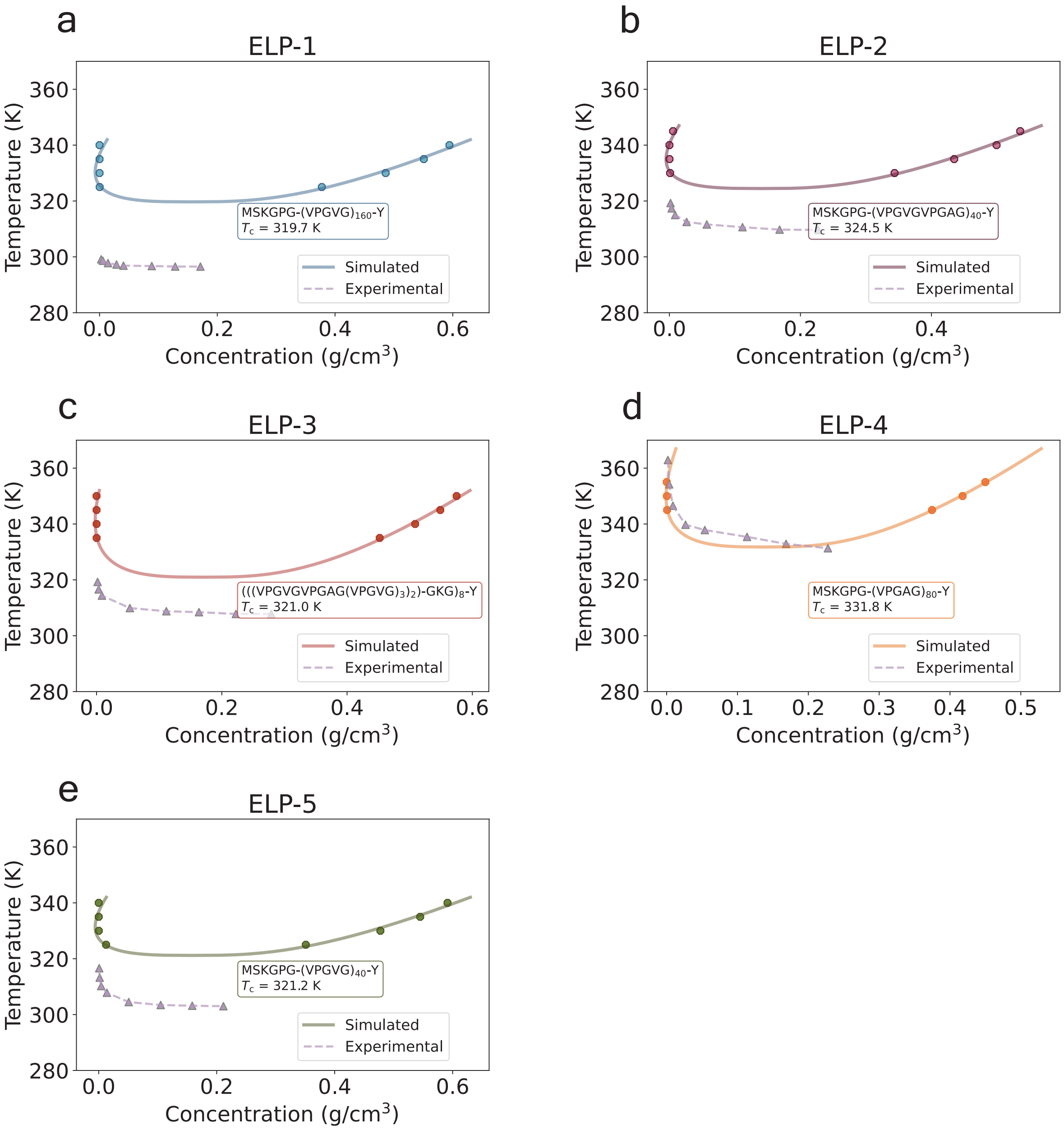


FIG. S3: **Mpipi-T Model 1 phase diagrams computed for:** (a) ELP-1, (b) ELP-2, (c) ELP-3, (d) ELP-4, and (e) ELP-5. The ELP sequence and the critical temperature extracted from simulations (using the law of coexistence densities and law of rectilinear diameters) are shown in the inset. Each trajectory is divided into three blocks, with the mean of the blocks used to determine each data point. The error bars, representing the standard error, are not shown as they are smaller than the size of the data points.

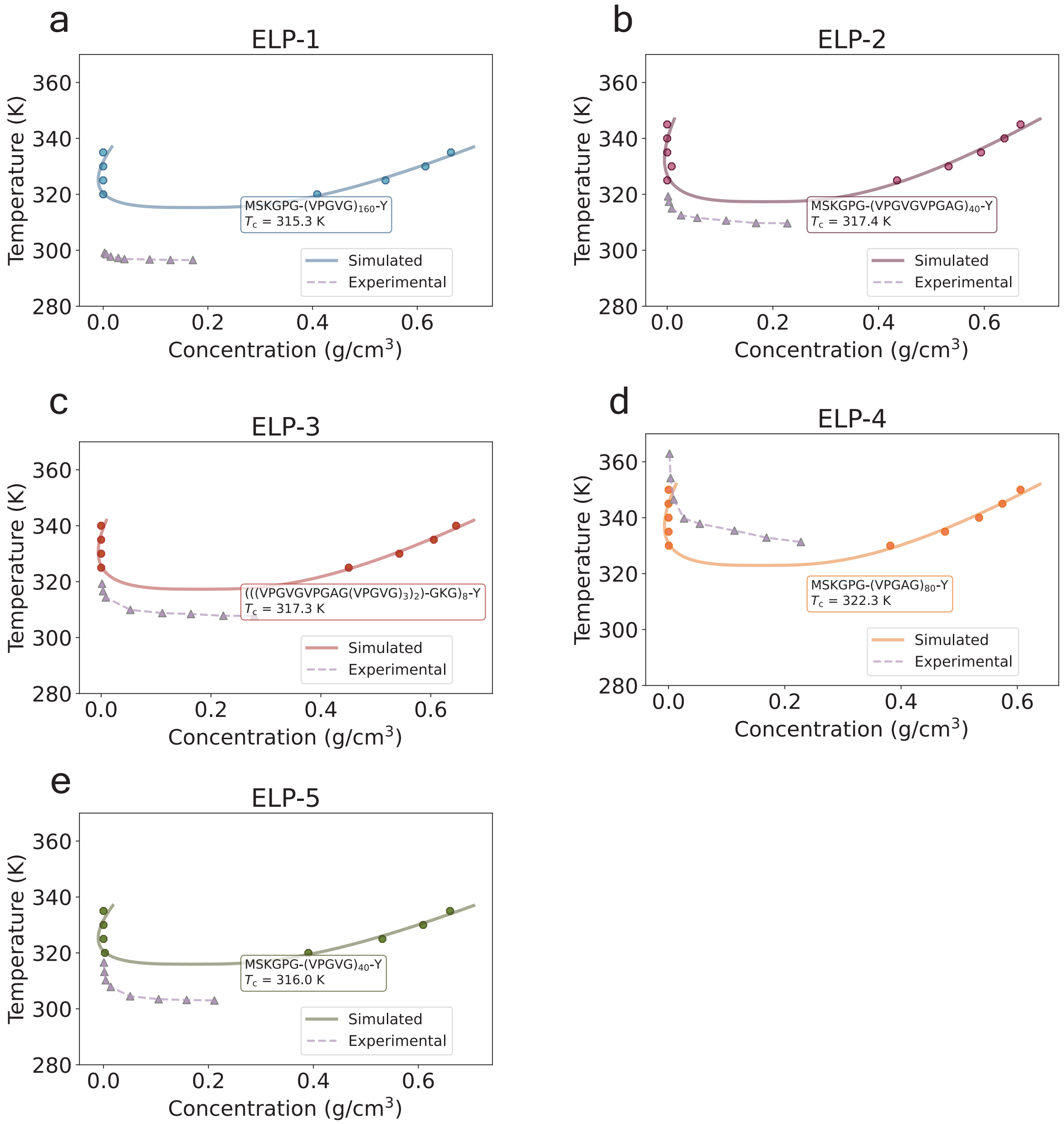


FIG. S4: **Mpipi-T Model 2 phase diagrams computed for:** (a) ELP-1, (b) ELP-2, (c) ELP-3, (d) ELP-4, and (e) ELP-5. The ELP sequence and the critical temperature extracted from simulations (using the law of coexistence densities and law of rectilinear diameters) are shown in the inset. Each trajectory is divided into three blocks, with the mean of the blocks used to determine each data point. The error bars, representing the standard error, are not shown as they are smaller than the size of the data points.

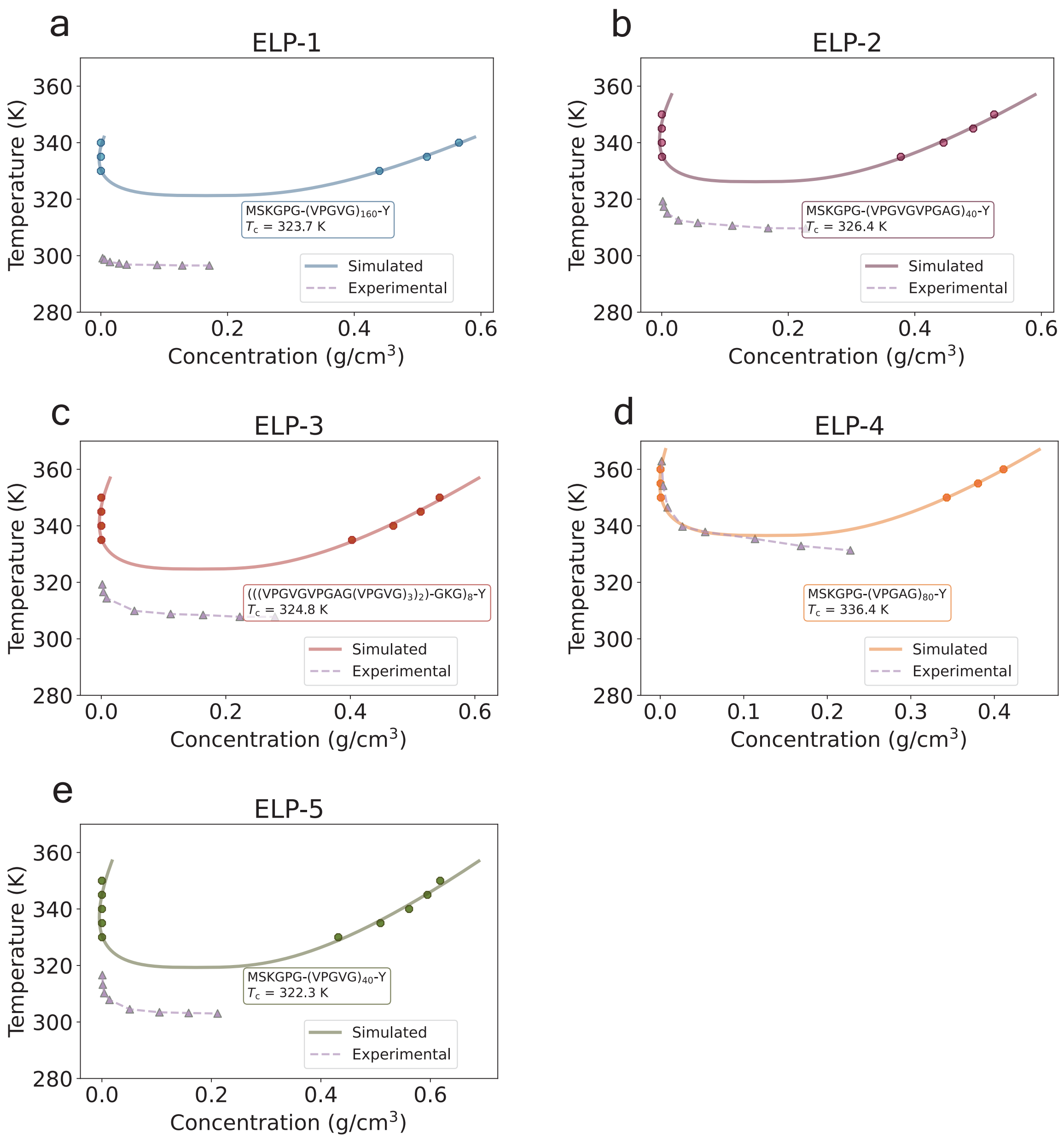


FIG. S5: **Mpipi-T Model 3 phase diagrams computed for:** (a) ELP-1, (b) ELP-2, (c) ELP-3, (d) ELP-4, and (e) ELP-5. The ELP sequence and the critical temperature extracted from simulations (using the law of coexistence densities and law of rectilinear diameters) are shown in the inset. Each trajectory is divided into three blocks, with the mean of the blocks used to determine each data point. The error bars, representing the standard error, are not shown as they are smaller than the size of the data points.

#### III. CLOUD-FIT: Cloud Point Computations

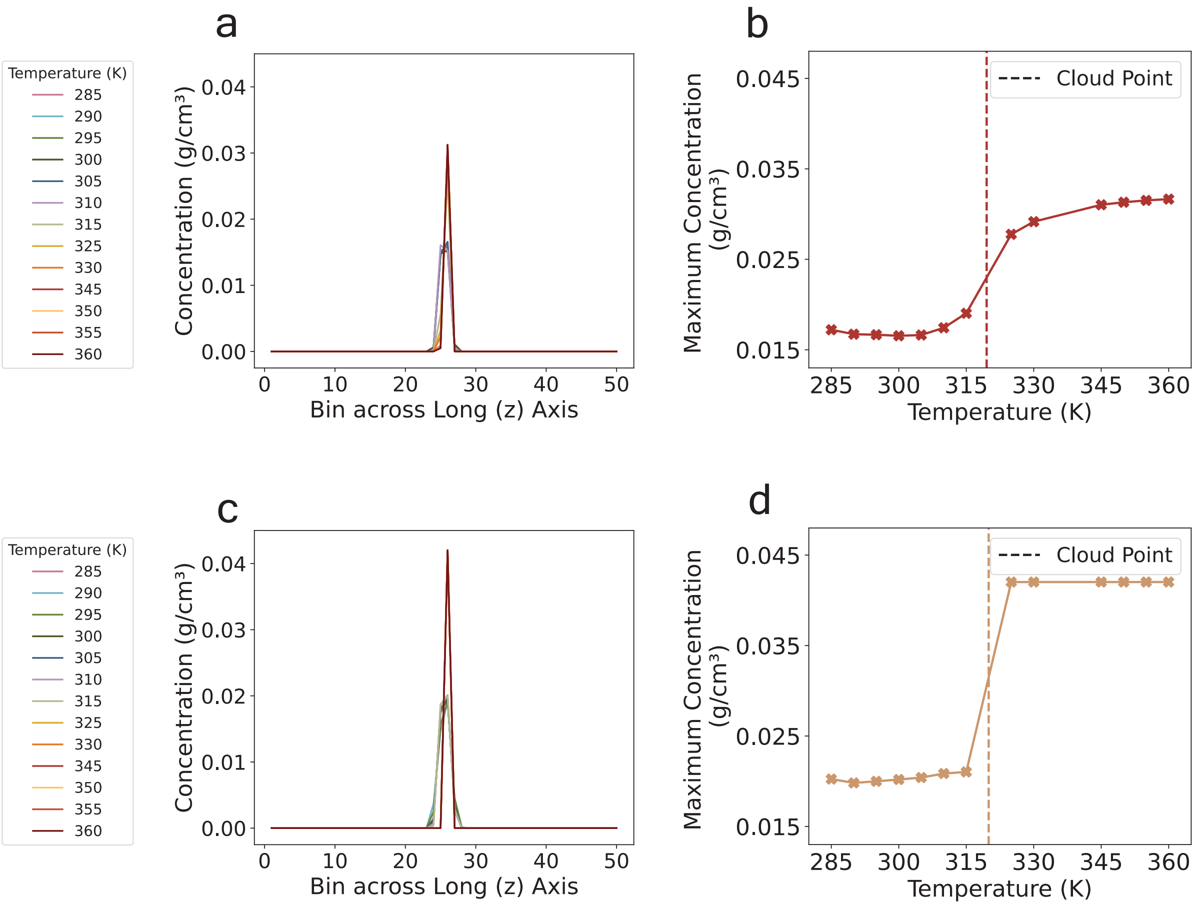


FIG. S6: **CLOUD-FIT: a new method to compute cloud point from simulations in the dilute phase is illustrated using two examples.** At a fixed system concentration, the cloud point is the temperature at which the system becomes turbid due to phase separation. In experiments, this is often measured using a low system concentration. For LCST sequences, the sample is heated gradually from a low system concentration until it reaches the left arm of the binodal. Since nucleation events are rare in dilute solutions, CLOUD-FIT exploits finite-size effects to capture density fluctuations at the target system concentration. A simulation box with proteins is prepared with 64 protein replicates (if protein length is less than 190 residues) or 27 replicates (if protein length is greater than or equal to 190 residues). The box is then compressed using *NPT* simulations to create a slab of high protein density, thereby accelerating the rate at which fluctuations are observed. The z-dimension (or long axis) is extended to achieve the target protein concentration. (a) and (c) show the the density profiles along the z-axis of the slab system after *NVT* simulations are performed for each temperature in small temperature intervals over the desired range. The system is scanned through, and the region of maximum concentration is recorded. The maximum concentration at this region for each of these temperatures in (a) and (c) is extracted and plotted in (b) and (d), respectively. At temperatures below the cloud point, the density is more distributed throughout the box, resulting in a lower maximum concentration. At temperatures above the cloud point, the system condenses, resulting in a higher maximum concentration. The midpoint of the region with the highest slope is determined to be the cloud point.

#### IV. Fitness Function Parameters for Model 3

Tables S2 to S21 are the fitness function parameters for $\varepsilon_{ij}$ in Model 3 of Mpipi-T. The parameters for Models 1 and 2 can be found in the Mpipi-T GitHub repository.

| Amino Acid $i$ | Amino Acid $j$ | $\varepsilon_{ii,\text{Mpipi}}$ | $\varepsilon_{jj,\text{Mpipi}}$ | $a_i$ | $b_i$ | $c_i$ | $\alpha_i$ | $a_j$ | $b_j$ | $c_j$ | $\alpha_j$ |
|---|---|---|---|---|---|---|---|---|---|---|---|
| A | R | 0.049480 | 0.089916 | -6.84807201e-05 | 5.42590026e-02 | -8.46961697e+00 | 0.70 | 0 | 0 | 0 | 0 |
| A | H | 0.049480 | 0.027216 | -6.84807201e-05 | 5.42590026e-02 | -8.46961697e+00 | 0.70 | 0 | 0 | 0 | 0 |
| A | K | 0.049480 | 0.019117 | -6.84807201e-05 | 5.42590026e-02 | -8.46961697e+00 | 0.70 | 0 | 0 | 0 | 0 |
| A | D | 0.049480 | 0.079096 | -6.84807201e-05 | 5.42590026e-02 | -8.46961697e+00 | 0.70 | 0 | 0 | 0 | 0 |
| A | E | 0.049480 | 0.085622 | -6.84807201e-05 | 5.42590026e-02 | -8.46961697e+00 | 0.70 | 0 | 0 | 0 | 0 |
| A | S | 0.049480 | 0.061600 | -6.84807201e-05 | 5.42590026e-02 | -8.46961697e+00 | 0.70 | 0 | 0 | 0 | 0 |
| A | T | 0.049480 | 0.030758 | -6.84807201e-05 | 5.42590026e-02 | -8.46961697e+00 | 0.70 | 0 | 0 | 0 | 0 |
| A | N | 0.049480 | 0.193849 | -6.84807201e-05 | 5.42590026e-02 | -8.46961697e+00 | 0.70 | 0 | 0 | 0 | 0 |
| A | Q | 0.049480 | 0.200448 | -6.84807201e-05 | 5.42590026e-02 | -8.46961697e+00 | 0.70 | 0 | 0 | 0 | 0 |
| A | C | 0.049480 | 0.069311 | -6.84807201e-05 | 5.42590026e-02 | -8.46961697e+00 | 0.70 | 0 | 0 | 0 | 0 |
| A | G | 0.049480 | 0.096470 | -1.09569152e-04 | 5.42590026e-02 | -8.46961697e+00 | 0.70 | 0 | 0 | 0 | 0 |
| A | P | 0.049480 | 0.078677 | -6.84807201e-05 | 5.42590026e-02 | -8.46961697e+00 | 0.70 | 0 | 0 | 0 | 0 |
| A | A | 0.049480 | 0.049480 | -6.84807201e-05 | 5.42590026e-02 | -8.46961697e+00 | 0.70 | -6.84807201e-05 | 5.42590026e-02 | -8.46961697e+00 | 0.70 |
| A | V | 0.049480 | 0.005578 | -6.84807201e-05 | 5.42590026e-02 | -8.46961697e+00 | 0.70 | -1.49763237e-04 | 1.15406051e-01 | -1.92322375e+01 | 0.70 |
| A | I | 0.049480 | 0.000395 | -6.84807201e-05 | 5.42590026e-02 | -8.46961697e+00 | 0.70 | -9.88175293e-05 | 1.10797972e-01 | -1.85218569e+01 | 0.70 |
| A | L | 0.049480 | 0.010998 | -6.84807201e-05 | 5.42590026e-02 | -8.46961697e+00 | 0.70 | -1.06970639e-04 | 1.17281979e-01 | -1.91871215e+01 | 0.70 |
| A | M | 0.049480 | 0.039564 | -6.84807201e-05 | 5.42590026e-02 | -8.46961697e+00 | 0.70 | -9.14334357e-05 | 8.01341277e-02 | -1.76908670e+01 | 0.70 |
| A | F | 0.049480 | 0.391642 | -6.84807201e-05 | 5.42590026e-02 | -8.46961697e+00 | 0.70 | 0 | 0 | 0 | 0 |
| A | Y | 0.049480 | 0.419186 | -6.84807201e-05 | 5.42590026e-02 | -8.46961697e+00 | 0.70 | 0 | 0 | 0 | 0 |
| A | W | 0.049480 | 0.550297 | -6.84807201e-05 | 5.42590026e-02 | -8.46961697e+00 | 0.70 | 0 | 0 | 0 | 0 |

TABLE S2: Fitness function parameters for $\varepsilon_{ij}$ where $i$ = A in Mpipi-T Model 3

| Amino Acid $i$ | Amino Acid $j$ | $\varepsilon_{ii,\text{Mpipi}}$ | $\varepsilon_{jj,\text{Mpipi}}$ | $a_i$ | $b_i$ | $c_i$ | $\alpha_i$ | $a_j$ | $b_j$ | $c_j$ | $\alpha_j$ |
|---|---|---|---|---|---|---|---|---|---|---|---|
| C | R | 0.069311 | 0.089916 | 0 | 0 | 0 | 0 | 0 | 0 | 0 | 0 |
| C | H | 0.069311 | 0.027216 | 0 | 0 | 0 | 0 | 0 | 0 | 0 | 0 |
| C | K | 0.069311 | 0.019117 | 0 | 0 | 0 | 0 | 0 | 0 | 0 | 0 |
| C | D | 0.069311 | 0.079096 | 0 | 0 | 0 | 0 | 0 | 0 | 0 | 0 |
| C | E | 0.069311 | 0.085622 | 0 | 0 | 0 | 0 | 0 | 0 | 0 | 0 |
| C | S | 0.069311 | 0.061600 | 0 | 0 | 0 | 0 | 0 | 0 | 0 | 0 |
| C | T | 0.069311 | 0.030758 | 0 | 0 | 0 | 0 | 0 | 0 | 0 | 0 |
| C | N | 0.069311 | 0.193849 | 0 | 0 | 0 | 0 | 0 | 0 | 0 | 0 |
| C | Q | 0.069311 | 0.200448 | 0 | 0 | 0 | 0 | 0 | 0 | 0 | 0 |
| C | C | 0.069311 | 0.069311 | 0 | 0 | 0 | 0 | 0 | 0 | 0 | 0 |
| C | G | 0.069311 | 0.096470 | 0 | 0 | 0 | 0 | 0 | 0 | 0 | 0 |
| C | P | 0.069311 | 0.078677 | 0 | 0 | 0 | 0 | 0 | 0 | 0 | 0 |
| C | A | 0.069311 | 0.049480 | 0 | 0 | 0 | 0 | -6.84807201e-05 | 5.42590026e-02 | -8.46961697e+00 | 0.70 |
| C | V | 0.069311 | 0.005578 | 0 | 0 | 0 | 0 | -1.49763237e-04 | 1.15406051e-01 | -1.92322375e+01 | 0.70 |
| C | I | 0.069311 | 0.000395 | 0 | 0 | 0 | 0 | -9.88175293e-05 | 1.10797972e-01 | -1.85218569e+01 | 0.70 |
| C | L | 0.069311 | 0.010998 | 0 | 0 | 0 | 0 | -1.06970639e-04 | 1.17281979e-01 | -1.91871215e+01 | 0.70 |
| C | M | 0.069311 | 0.039564 | 0 | 0 | 0 | 0 | -9.14334357e-05 | 8.01341277e-02 | -1.76908670e+01 | 0.70 |
| C | F | 0.069311 | 0.391642 | 0 | 0 | 0 | 0 | 0 | 0 | 0 | 0 |
| C | Y | 0.069311 | 0.419186 | 0 | 0 | 0 | 0 | 0 | 0 | 0 | 0 |
| C | W | 0.069311 | 0.550297 | 0 | 0 | 0 | 0 | 0 | 0 | 0 | 0 |

TABLE S3: Fitness function parameters for $\varepsilon_{ij}$ where $i$ = C in Mpipi-T Model 3

| Amino Acid $i$ | Amino Acid $j$ | $\varepsilon_{ii,\text{Mpipi}}$ | $\varepsilon_{jj,\text{Mpipi}}$ | $a_i$ | $b_i$ | $c_i$ | $\alpha_i$ | $a_j$ | $b_j$ | $c_j$ | $\alpha_j$ |
|---|---|---|---|---|---|---|---|---|---|---|---|
| D | R | 0.079096 | 0.089916 | 0 | 0 | 0 | 0 | 0 | 0 | 0 | 0 |
| D | H | 0.079096 | 0.027216 | 0 | 0 | 0 | 0 | 0 | 0 | 0 | 0 |
| D | K | 0.079096 | 0.019117 | 0 | 0 | 0 | 0 | 0 | 0 | 0 | 0 |
| D | D | 0.079096 | 0.079096 | 0 | 0 | 0 | 0 | 0 | 0 | 0 | 0 |
| D | E | 0.079096 | 0.085622 | 0 | 0 | 0 | 0 | 0 | 0 | 0 | 0 |
| D | S | 0.079096 | 0.061600 | 0 | 0 | 0 | 0 | 0 | 0 | 0 | 0 |
| D | T | 0.079096 | 0.030758 | 0 | 0 | 0 | 0 | 0 | 0 | 0 | 0 |
| D | N | 0.079096 | 0.193849 | 0 | 0 | 0 | 0 | 0 | 0 | 0 | 0 |
| D | Q | 0.079096 | 0.200448 | 0 | 0 | 0 | 0 | 0 | 0 | 0 | 0 |
| D | C | 0.079096 | 0.069311 | 0 | 0 | 0 | 0 | 0 | 0 | 0 | 0 |
| D | G | 0.079096 | 0.096470 | 0 | 0 | 0 | 0 | 0 | 0 | 0 | 0 |
| D | P | 0.079096 | 0.078677 | 0 | 0 | 0 | 0 | 0 | 0 | 0 | 0 |
| D | A | 0.079096 | 0.049480 | 0 | 0 | 0 | 0 | -6.84807201e-05 | 5.42590026e-02 | -8.46961697e+00 | 0.70 |
| D | V | 0.079096 | 0.005578 | 0 | 0 | 0 | 0 | -1.49763237e-04 | 1.15406051e-01 | -1.92322375e+01 | 0.70 |
| D | I | 0.079096 | 0.000395 | 0 | 0 | 0 | 0 | -9.88175293e-05 | 1.10797972e-01 | -1.85218569e+01 | 0.70 |
| D | L | 0.079096 | 0.010998 | 0 | 0 | 0 | 0 | -1.06970639e-04 | 1.17281979e-01 | -1.91871215e+01 | 0.70 |
| D | M | 0.079096 | 0.039564 | 0 | 0 | 0 | 0 | -9.14334357e-05 | 8.01341277e-02 | -1.76908670e+01 | 0.70 |
| D | F | 0.079096 | 0.391642 | 0 | 0 | 0 | 0 | 0 | 0 | 0 | 0 |
| D | Y | 0.079096 | 0.419186 | 0 | 0 | 0 | 0 | 0 | 0 | 0 | 0 |
| D | W | 0.079096 | 0.550297 | 0 | 0 | 0 | 0 | 0 | 0 | 0 | 0 |

TABLE S4: Fitness function parameters for $\varepsilon_{ij}$ where $i$ = D in Mpipi-T Model 3

| Amino Acid $i$ | Amino Acid $j$ | $\varepsilon_{ii,\text{Mpipi}}$ | $\varepsilon_{jj,\text{Mpipi}}$ | $a_i$ | $b_i$ | $c_i$ | $\alpha_i$ | $a_j$ | $b_j$ | $c_j$ | $\alpha_j$ |
|---|---|---|---|---|---|---|---|---|---|---|---|
| E | R | 0.085622 | 0.089916 | 0 | 0 | 0 | 0 | 0 | 0 | 0 | 0 |
| E | H | 0.085622 | 0.027216 | 0 | 0 | 0 | 0 | 0 | 0 | 0 | 0 |
| E | K | 0.085622 | 0.019117 | 0 | 0 | 0 | 0 | 0 | 0 | 0 | 0 |
| E | D | 0.085622 | 0.079096 | 0 | 0 | 0 | 0 | 0 | 0 | 0 | 0 |
| E | E | 0.085622 | 0.085622 | 0 | 0 | 0 | 0 | 0 | 0 | 0 | 0 |
| E | S | 0.085622 | 0.061600 | 0 | 0 | 0 | 0 | 0 | 0 | 0 | 0 |
| E | T | 0.085622 | 0.030758 | 0 | 0 | 0 | 0 | 0 | 0 | 0 | 0 |
| E | N | 0.085622 | 0.193849 | 0 | 0 | 0 | 0 | 0 | 0 | 0 | 0 |
| E | Q | 0.085622 | 0.200448 | 0 | 0 | 0 | 0 | 0 | 0 | 0 | 0 |
| E | C | 0.085622 | 0.069311 | 0 | 0 | 0 | 0 | 0 | 0 | 0 | 0 |
| E | G | 0.085622 | 0.096470 | 0 | 0 | 0 | 0 | 0 | 0 | 0 | 0 |
| E | P | 0.085622 | 0.078677 | 0 | 0 | 0 | 0 | 0 | 0 | 0 | 0 |
| E | A | 0.085622 | 0.049480 | 0 | 0 | 0 | 0 | -6.84807201e-05 | 5.42590026e-02 | -8.46961697e+00 | 0.70 |
| E | V | 0.085622 | 0.005578 | 0 | 0 | 0 | 0 | -1.49763237e-04 | 1.15406051e-01 | -1.92322375e+01 | 0.70 |
| E | I | 0.085622 | 0.000395 | 0 | 0 | 0 | 0 | -9.88175293e-05 | 1.10797972e-01 | -1.85218569e+01 | 0.70 |
| E | L | 0.085622 | 0.010998 | 0 | 0 | 0 | 0 | -1.06970639e-04 | 1.17281979e-01 | -1.91871215e+01 | 0.70 |
| E | M | 0.085622 | 0.039564 | 0 | 0 | 0 | 0 | -9.14334357e-05 | 8.01341277e-02 | -1.76908670e+01 | 0.70 |
| E | F | 0.085622 | 0.391642 | 0 | 0 | 0 | 0 | 0 | 0 | 0 | 0 |
| E | Y | 0.085622 | 0.419186 | 0 | 0 | 0 | 0 | 0 | 0 | 0 | 0 |
| E | W | 0.085622 | 0.550297 | 0 | 0 | 0 | 0 | 0 | 0 | 0 | 0 |

TABLE S5: Fitness function parameters for $\varepsilon_{ij}$ where $i$ = E in Mpipi-T Model 3

| Amino Acid $i$ | Amino Acid $j$ | $\varepsilon_{ii,\text{Mpipi}}$ | $\varepsilon_{jj,\text{Mpipi}}$ | $a_i$ | $b_i$ | $c_i$ | $\alpha_i$ | $a_j$ | $b_j$ | $c_j$ | $\alpha_j$ |
|---|---|---|---|---|---|---|---|---|---|---|---|
| F | R | 0.391642 | 0.089916 | 0 | 0 | 0 | 0 | 0 | 0 | 0 | 0 |
| F | H | 0.391642 | 0.027216 | 0 | 0 | 0 | 0 | 0 | 0 | 0 | 0 |
| F | K | 0.391642 | 0.019117 | 0 | 0 | 0 | 0 | 0 | 0 | 0 | 0 |
| F | D | 0.391642 | 0.079096 | 0 | 0 | 0 | 0 | 0 | 0 | 0 | 0 |
| F | E | 0.391642 | 0.085622 | 0 | 0 | 0 | 0 | 0 | 0 | 0 | 0 |
| F | S | 0.391642 | 0.061600 | 0 | 0 | 0 | 0 | 0 | 0 | 0 | 0 |
| F | T | 0.391642 | 0.030758 | 0 | 0 | 0 | 0 | 0 | 0 | 0 | 0 |
| F | N | 0.391642 | 0.193849 | 0 | 0 | 0 | 0 | 0 | 0 | 0 | 0 |
| F | Q | 0.391642 | 0.200448 | 0 | 0 | 0 | 0 | 0 | 0 | 0 | 0 |
| F | C | 0.391642 | 0.069311 | 0 | 0 | 0 | 0 | 0 | 0 | 0 | 0 |
| F | G | 0.391642 | 0.096470 | 0 | 0 | 0 | 0 | 0 | 0 | 0 | 0 |
| F | P | 0.391642 | 0.078677 | 0 | 0 | 0 | 0 | 0 | 0 | 0 | 0 |
| F | A | 0.391642 | 0.049480 | 0 | 0 | 0 | 0 | -6.84807201e-05 | 5.42590026e-02 | -8.46961697e+00 | 0.70 |
| F | V | 0.391642 | 0.005578 | 0 | 0 | 0 | 0 | -1.49763237e-04 | 1.15406051e-01 | -1.92322375e+01 | 0.70 |
| F | I | 0.391642 | 0.000395 | 0 | 0 | 0 | 0 | -9.88175293e-05 | 1.10797972e-01 | -1.85218569e+01 | 0.70 |
| F | L | 0.391642 | 0.010998 | 0 | 0 | 0 | 0 | -1.06970639e-04 | 1.17281979e-01 | -1.91871215e+01 | 0.70 |
| F | M | 0.391642 | 0.039564 | 0 | 0 | 0 | 0 | -9.14334357e-05 | 8.01341277e-02 | -1.76908670e+01 | 0.70 |
| F | F | 0.391642 | 0.391642 | 0 | 0 | 0 | 0 | 0 | 0 | 0 | 0 |
| F | Y | 0.391642 | 0.419186 | 0 | 0 | 0 | 0 | 0 | 0 | 0 | 0 |
| F | W | 0.391642 | 0.550297 | 0 | 0 | 0 | 0 | 0 | 0 | 0 | 0 |

TABLE S6: Fitness function parameters for $\varepsilon_{ij}$ where $i$ = F in Mpipi-T Model 3

| Amino Acid $i$ | Amino Acid $j$ | $\varepsilon_{ii,\text{Mpipi}}$ | $\varepsilon_{jj,\text{Mpipi}}$ | $a_i$ | $b_i$ | $c_i$ | $\alpha_i$ | $a_j$ | $b_j$ | $c_j$ | $\alpha_j$ |
|---|---|---|---|---|---|---|---|---|---|---|---|
| G | R | 0.096470 | 0.089916 | 0 | 0 | 0 | 0 | 0 | 0 | 0 | 0 |
| G | H | 0.096470 | 0.027216 | 0 | 0 | 0 | 0 | 0 | 0 | 0 | 0 |
| G | K | 0.096470 | 0.019117 | 0 | 0 | 0 | 0 | 0 | 0 | 0 | 0 |
| G | D | 0.096470 | 0.079096 | 0 | 0 | 0 | 0 | 0 | 0 | 0 | 0 |
| G | E | 0.096470 | 0.085622 | 0 | 0 | 0 | 0 | 0 | 0 | 0 | 0 |
| G | S | 0.096470 | 0.061600 | 0 | 0 | 0 | 0 | 0 | 0 | 0 | 0 |
| G | T | 0.096470 | 0.030758 | 0 | 0 | 0 | 0 | 0 | 0 | 0 | 0 |
| G | N | 0.096470 | 0.193849 | 0 | 0 | 0 | 0 | 0 | 0 | 0 | 0 |
| G | Q | 0.096470 | 0.200448 | 0 | 0 | 0 | 0 | 0 | 0 | 0 | 0 |
| G | C | 0.096470 | 0.069311 | 0 | 0 | 0 | 0 | 0 | 0 | 0 | 0 |
| G | G | 0.096470 | 0.096470 | 0 | 0 | 0 | 0 | 0 | 0 | 0 | 0 |
| G | P | 0.096470 | 0.078677 | 0 | 0 | 0 | 0 | 0 | 0 | 0 | 0 |
| G | A | 0.096470 | 0.049480 | 0 | 0 | 0 | 0 | -1.09569152e-04 | 5.42590026e-02 | -8.46961697e+00 | 0.70 |
| G | V | 0.096470 | 0.005578 | 0 | 0 | 0 | 0 | -2.69573827e-04 | 1.15406051e-01 | -1.92322375e+01 | 0.70 |
| G | I | 0.096470 | 0.000395 | 0 | 0 | 0 | 0 | -1.77871553e-04 | 1.10797972e-01 | -1.85218569e+01 | 0.70 |
| G | L | 0.096470 | 0.010998 | 0 | 0 | 0 | 0 | -1.92547149e-04 | 1.17281979e-01 | -1.91871215e+01 | 0.70 |
| G | M | 0.096470 | 0.039564 | 0 | 0 | 0 | 0 | -1.64580184e-04 | 8.01341277e-02 | -1.76908670e+01 | 0.70 |
| G | F | 0.096470 | 0.391642 | 0 | 0 | 0 | 0 | 0 | 0 | 0 | 0 |
| G | Y | 0.096470 | 0.419186 | 0 | 0 | 0 | 0 | 0 | 0 | 0 | 0 |
| G | W | 0.096470 | 0.550297 | 0 | 0 | 0 | 0 | 0 | 0 | 0 | 0 |

TABLE S7: Fitness function parameters for $\varepsilon_{ij}$ where $i$ = G in Mpipi-T Model 3

| Amino Acid $i$ | Amino Acid $j$ | $\varepsilon_{ii,\text{Mpipi}}$ | $\varepsilon_{jj,\text{Mpipi}}$ | $a_i$ | $b_i$ | $c_i$ | $\alpha_i$ | $a_j$ | $b_j$ | $c_j$ | $\alpha_j$ |
|---|---|---|---|---|---|---|---|---|---|---|---|
| H | R | 0.027216 | 0.089916 | 0 | 0 | 0 | 0 | 0 | 0 | 0 | 0 |
| H | H | 0.027216 | 0.027216 | 0 | 0 | 0 | 0 | 0 | 0 | 0 | 0 |
| H | K | 0.027216 | 0.019117 | 0 | 0 | 0 | 0 | 0 | 0 | 0 | 0 |
| H | D | 0.027216 | 0.079096 | 0 | 0 | 0 | 0 | 0 | 0 | 0 | 0 |
| H | E | 0.027216 | 0.085622 | 0 | 0 | 0 | 0 | 0 | 0 | 0 | 0 |
| H | S | 0.027216 | 0.061600 | 0 | 0 | 0 | 0 | 0 | 0 | 0 | 0 |
| H | T | 0.027216 | 0.030758 | 0 | 0 | 0 | 0 | 0 | 0 | 0 | 0 |
| H | N | 0.027216 | 0.193849 | 0 | 0 | 0 | 0 | 0 | 0 | 0 | 0 |
| H | Q | 0.027216 | 0.200448 | 0 | 0 | 0 | 0 | 0 | 0 | 0 | 0 |
| H | C | 0.027216 | 0.069311 | 0 | 0 | 0 | 0 | 0 | 0 | 0 | 0 |
| H | G | 0.027216 | 0.096470 | 0 | 0 | 0 | 0 | 0 | 0 | 0 | 0 |
| H | P | 0.027216 | 0.078677 | 0 | 0 | 0 | 0 | 0 | 0 | 0 | 0 |
| H | A | 0.027216 | 0.049480 | 0 | 0 | 0 | 0 | -6.84807201e-05 | 5.42590026e-02 | -8.46961697e+00 | 0.70 |
| H | V | 0.027216 | 0.005578 | 0 | 0 | 0 | 0 | -1.49763237e-04 | 1.15406051e-01 | -1.92322375e+01 | 0.70 |
| H | I | 0.027216 | 0.000395 | 0 | 0 | 0 | 0 | -9.88175293e-05 | 1.10797972e-01 | -1.85218569e+01 | 0.70 |
| H | L | 0.027216 | 0.010998 | 0 | 0 | 0 | 0 | -1.06970639e-04 | 1.17281979e-01 | -1.91871215e+01 | 0.70 |
| H | M | 0.027216 | 0.039564 | 0 | 0 | 0 | 0 | -9.14334357e-05 | 8.01341277e-02 | -1.76908670e+01 | 0.70 |
| H | F | 0.027216 | 0.391642 | 0 | 0 | 0 | 0 | 0 | 0 | 0 | 0 |
| H | Y | 0.027216 | 0.419186 | 0 | 0 | 0 | 0 | 0 | 0 | 0 | 0 |
| H | W | 0.027216 | 0.550297 | 0 | 0 | 0 | 0 | 0 | 0 | 0 | 0 |

TABLE S8: Fitness function parameters for $\varepsilon_{ij}$ where $i$ = H in Mpipi-T Model 3

| Amino Acid $i$ | Amino Acid $j$ | $\varepsilon_{ii,\text{Mpipi}}$ | $\varepsilon_{jj,\text{Mpipi}}$ | $a_i$ | $b_i$ | $c_i$ | $\alpha_i$ | $a_j$ | $b_j$ | $c_j$ | $\alpha_j$ |
|---|---|---|---|---|---|---|---|---|---|---|---|
| I | R | 0.000395 | 0.089916 | -9.88175293e-05 | 1.10797972e-01 | -1.85218569e+01 | 0.70 | 0 | 0 | 0 | 0 |
| I | H | 0.000395 | 0.027216 | -9.88175293e-05 | 1.10797972e-01 | -1.85218569e+01 | 0.70 | 0 | 0 | 0 | 0 |
| I | K | 0.000395 | 0.019117 | -9.88175293e-05 | 1.10797972e-01 | -1.85218569e+01 | 0.70 | 0 | 0 | 0 | 0 |
| I | D | 0.000395 | 0.079096 | -9.88175293e-05 | 1.10797972e-01 | -1.85218569e+01 | 0.70 | 0 | 0 | 0 | 0 |
| I | E | 0.000395 | 0.085622 | -9.88175293e-05 | 1.10797972e-01 | -1.85218569e+01 | 0.70 | 0 | 0 | 0 | 0 |
| I | S | 0.000395 | 0.061600 | -9.88175293e-05 | 1.10797972e-01 | -1.85218569e+01 | 0.70 | 0 | 0 | 0 | 0 |
| I | T | 0.000395 | 0.030758 | -9.88175293e-05 | 1.10797972e-01 | -1.85218569e+01 | 0.70 | 0 | 0 | 0 | 0 |
| I | N | 0.000395 | 0.193849 | -9.88175293e-05 | 1.10797972e-01 | -1.85218569e+01 | 0.70 | 0 | 0 | 0 | 0 |
| I | Q | 0.000395 | 0.200448 | -9.88175293e-05 | 1.10797972e-01 | -1.85218569e+01 | 0.70 | 0 | 0 | 0 | 0 |
| I | C | 0.000395 | 0.069311 | -9.88175293e-05 | 1.10797972e-01 | -1.85218569e+01 | 0.70 | 0 | 0 | 0 | 0 |
| I | G | 0.000395 | 0.096470 | -1.77871553e-04 | 1.10797972e-01 | -1.85218569e+01 | 0.70 | 0 | 0 | 0 | 0 |
| I | P | 0.000395 | 0.078677 | -9.88175293e-05 | 1.10797972e-01 | -1.85218569e+01 | 0.70 | 0 | 0 | 0 | 0 |
| I | A | 0.000395 | 0.049480 | -9.88175293e-05 | 1.10797972e-01 | -1.85218569e+01 | 0.70 | -6.84807201e-05 | 5.42590026e-02 | -8.46961697e+00 | 0.70 |
| I | V | 0.000395 | 0.005578 | -9.88175293e-05 | 1.10797972e-01 | -1.85218569e+01 | 0.70 | -1.49763237e-04 | 1.15406051e-01 | -1.92322375e+01 | 0.70 |
| I | I | 0.000395 | 0.000395 | -9.88175293e-05 | 1.10797972e-01 | -1.85218569e+01 | 0.70 | -9.88175293e-05 | 1.10797972e-01 | -1.85218569e+01 | 0.70 |
| I | L | 0.000395 | 0.010998 | -9.88175293e-05 | 1.10797972e-01 | -1.85218569e+01 | 0.70 | -1.06970639e-04 | 1.17281979e-01 | -1.91871215e+01 | 0.70 |
| I | M | 0.000395 | 0.039564 | -9.88175293e-05 | 1.10797972e-01 | -1.85218569e+01 | 0.70 | -9.14334357e-05 | 8.01341277e-02 | -1.76908670e+01 | 0.70 |
| I | F | 0.000395 | 0.391642 | -9.88175293e-05 | 1.10797972e-01 | -1.85218569e+01 | 0.70 | 0 | 0 | 0 | 0 |
| I | Y | 0.000395 | 0.419186 | -9.88175293e-05 | 1.10797972e-01 | -1.85218569e+01 | 0.70 | 0 | 0 | 0 | 0 |
| I | W | 0.000395 | 0.550297 | -9.88175293e-05 | 1.10797972e-01 | -1.85218569e+01 | 0.70 | 0 | 0 | 0 | 0 |

TABLE S9: Fitness function parameters for $\varepsilon_{ij}$ where $i$ = I in Mpipi-T Model 3

| Amino Acid $i$ | Amino Acid $j$ | $\varepsilon_{ii,\text{Mpipi}}$ | $\varepsilon_{jj,\text{Mpipi}}$ | $a_i$ | $b_i$ | $c_i$ | $\alpha_i$ | $a_j$ | $b_j$ | $c_j$ | $\alpha_j$ |
|---|---|---|---|---|---|---|---|---|---|---|---|
| K | R | 0.019117 | 0.089916 | 0 | 0 | 0 | 0 | 0 | 0 | 0 | 0 |
| K | H | 0.019117 | 0.027216 | 0 | 0 | 0 | 0 | 0 | 0 | 0 | 0 |
| K | K | 0.019117 | 0.019117 | 0 | 0 | 0 | 0 | 0 | 0 | 0 | 0 |
| K | D | 0.019117 | 0.079096 | 0 | 0 | 0 | 0 | 0 | 0 | 0 | 0 |
| K | E | 0.019117 | 0.085622 | 0 | 0 | 0 | 0 | 0 | 0 | 0 | 0 |
| K | S | 0.019117 | 0.061600 | 0 | 0 | 0 | 0 | 0 | 0 | 0 | 0 |
| K | T | 0.019117 | 0.030758 | 0 | 0 | 0 | 0 | 0 | 0 | 0 | 0 |
| K | N | 0.019117 | 0.193849 | 0 | 0 | 0 | 0 | 0 | 0 | 0 | 0 |
| K | Q | 0.019117 | 0.200448 | 0 | 0 | 0 | 0 | 0 | 0 | 0 | 0 |
| K | C | 0.019117 | 0.069311 | 0 | 0 | 0 | 0 | 0 | 0 | 0 | 0 |
| K | G | 0.019117 | 0.096470 | 0 | 0 | 0 | 0 | 0 | 0 | 0 | 0 |
| K | P | 0.019117 | 0.078677 | 0 | 0 | 0 | 0 | 0 | 0 | 0 | 0 |
| K | A | 0.019117 | 0.049480 | 0 | 0 | 0 | 0 | -6.84807201e-05 | 5.42590026e-02 | -8.46961697e+00 | 0.70 |
| K | V | 0.019117 | 0.005578 | 0 | 0 | 0 | 0 | -1.49763237e-04 | 1.15406051e-01 | -1.92322375e+01 | 0.70 |
| K | I | 0.019117 | 0.000395 | 0 | 0 | 0 | 0 | -9.88175293e-05 | 1.10797972e-01 | -1.85218569e+01 | 0.70 |
| K | L | 0.019117 | 0.010998 | 0 | 0 | 0 | 0 | -1.06970639e-04 | 1.17281979e-01 | -1.91871215e+01 | 0.70 |
| K | M | 0.019117 | 0.039564 | 0 | 0 | 0 | 0 | -9.14334357e-05 | 8.01341277e-02 | -1.76908670e+01 | 0.70 |
| K | F | 0.019117 | 0.391642 | 0 | 0 | 0 | 0 | 0 | 0 | 0 | 0 |
| K | Y | 0.019117 | 0.419186 | 0 | 0 | 0 | 0 | 0 | 0 | 0 | 0 |
| K | W | 0.019117 | 0.550297 | 0 | 0 | 0 | 0 | 0 | 0 | 0 | 0 |

TABLE S10: Fitness function parameters for $\varepsilon_{ij}$ where $i$ = K in Mpipi-T Model 3

| Amino Acid $i$ | Amino Acid $j$ | $\varepsilon_{ii,\text{Mpipi}}$ | $\varepsilon_{jj,\text{Mpipi}}$ | $a_i$ | $b_i$ | $c_i$ | $\alpha_i$ | $a_j$ | $b_j$ | $c_j$ | $\alpha_j$ |
|---|---|---|---|---|---|---|---|---|---|---|---|
| L | R | 0.010998 | 0.089916 | -1.06970639e-04 | 1.17281979e-01 | -1.91871215e+01 | 0.70 | 0 | 0 | 0 | 0 |
| L | H | 0.010998 | 0.027216 | -1.06970639e-04 | 1.17281979e-01 | -1.91871215e+01 | 0.70 | 0 | 0 | 0 | 0 |
| L | K | 0.010998 | 0.019117 | -1.06970639e-04 | 1.17281979e-01 | -1.91871215e+01 | 0.70 | 0 | 0 | 0 | 0 |
| L | D | 0.010998 | 0.079096 | -1.06970639e-04 | 1.17281979e-01 | -1.91871215e+01 | 0.70 | 0 | 0 | 0 | 0 |
| L | E | 0.010998 | 0.085622 | -1.06970639e-04 | 1.17281979e-01 | -1.91871215e+01 | 0.70 | 0 | 0 | 0 | 0 |
| L | S | 0.010998 | 0.061600 | -1.06970639e-04 | 1.17281979e-01 | -1.91871215e+01 | 0.70 | 0 | 0 | 0 | 0 |
| L | T | 0.010998 | 0.030758 | -1.06970639e-04 | 1.17281979e-01 | -1.91871215e+01 | 0.70 | 0 | 0 | 0 | 0 |
| L | N | 0.010998 | 0.193849 | -1.06970639e-04 | 1.17281979e-01 | -1.91871215e+01 | 0.70 | 0 | 0 | 0 | 0 |
| L | Q | 0.010998 | 0.200448 | -1.06970639e-04 | 1.17281979e-01 | -1.91871215e+01 | 0.70 | 0 | 0 | 0 | 0 |
| L | C | 0.010998 | 0.069311 | -1.06970639e-04 | 1.17281979e-01 | -1.91871215e+01 | 0.70 | 0 | 0 | 0 | 0 |
| L | G | 0.010998 | 0.096470 | -1.92547149e-04 | 1.17281979e-01 | -1.91871215e+01 | 0.70 | 0 | 0 | 0 | 0 |
| L | P | 0.010998 | 0.078677 | -1.06970639e-04 | 1.17281979e-01 | -1.91871215e+01 | 0.70 | 0 | 0 | 0 | 0 |
| L | A | 0.010998 | 0.049480 | -1.06970639e-04 | 1.17281979e-01 | -1.91871215e+01 | 0.70 | -6.84807201e-05 | 5.42590026e-02 | -8.46961697e+00 | 0.70 |
| L | V | 0.010998 | 0.005578 | -1.06970639e-04 | 1.17281979e-01 | -1.91871215e+01 | 0.70 | -1.49763237e-04 | 1.15406051e-01 | -1.92322375e+01 | 0.70 |
| L | I | 0.010998 | 0.000395 | -1.06970639e-04 | 1.17281979e-01 | -1.91871215e+01 | 0.70 | -9.88175293e-05 | 1.10797972e-01 | -1.85218569e+01 | 0.70 |
| L | L | 0.010998 | 0.010998 | -1.06970639e-04 | 1.17281979e-01 | -1.91871215e+01 | 0.70 | -1.06970639e-04 | 1.17281979e-01 | -1.91871215e+01 | 0.70 |
| L | M | 0.010998 | 0.039564 | -1.06970639e-04 | 1.17281979e-01 | -1.91871215e+01 | 0.70 | -9.14334357e-05 | 8.01341277e-02 | -1.76908670e+01 | 0.70 |
| L | F | 0.010998 | 0.391642 | -1.06970639e-04 | 1.17281979e-01 | -1.91871215e+01 | 0.70 | 0 | 0 | 0 | 0 |
| L | Y | 0.010998 | 0.419186 | -1.06970639e-04 | 1.17281979e-01 | -1.91871215e+01 | 0.70 | 0 | 0 | 0 | 0 |
| L | W | 0.010998 | 0.550297 | -1.06970639e-04 | 1.17281979e-01 | -1.91871215e+01 | 0.70 | 0 | 0 | 0 | 0 |

TABLE S11: Fitness function parameters for $\varepsilon_{ij}$ where $i$ = L in Mpipi-T Model 3

| Amino Acid $i$ | Amino Acid $j$ | $\varepsilon_{ii,\text{Mpipi}}$ | $\varepsilon_{jj,\text{Mpipi}}$ | $a_i$ | $b_i$ | $c_i$ | $\alpha_i$ | $a_j$ | $b_j$ | $c_j$ | $\alpha_j$ |
|---|---|---|---|---|---|---|---|---|---|---|---|
| M | R | 0.039564 | 0.089916 | -9.14334357e-05 | 8.01341277e-02 | -1.76908670e+01 | 0.70 | 0 | 0 | 0 | 0 |
| M | H | 0.039564 | 0.027216 | -9.14334357e-05 | 8.01341277e-02 | -1.76908670e+01 | 0.70 | 0 | 0 | 0 | 0 |
| M | K | 0.039564 | 0.019117 | -9.14334357e-05 | 8.01341277e-02 | -1.76908670e+01 | 0.70 | 0 | 0 | 0 | 0 |
| M | D | 0.039564 | 0.079096 | -9.14334357e-05 | 8.01341277e-02 | -1.76908670e+01 | 0.70 | 0 | 0 | 0 | 0 |
| M | E | 0.039564 | 0.085622 | -9.14334357e-05 | 8.01341277e-02 | -1.76908670e+01 | 0.70 | 0 | 0 | 0 | 0 |
| M | S | 0.039564 | 0.061600 | -9.14334357e-05 | 8.01341277e-02 | -1.76908670e+01 | 0.70 | 0 | 0 | 0 | 0 |
| M | T | 0.039564 | 0.030758 | -9.14334357e-05 | 8.01341277e-02 | -1.76908670e+01 | 0.70 | 0 | 0 | 0 | 0 |
| M | N | 0.039564 | 0.193849 | -9.14334357e-05 | 8.01341277e-02 | -1.76908670e+01 | 0.70 | 0 | 0 | 0 | 0 |
| M | Q | 0.039564 | 0.200448 | -9.14334357e-05 | 8.01341277e-02 | -1.76908670e+01 | 0.70 | 0 | 0 | 0 | 0 |
| M | C | 0.039564 | 0.069311 | -9.14334357e-05 | 8.01341277e-02 | -1.76908670e+01 | 0.70 | 0 | 0 | 0 | 0 |
| M | G | 0.039564 | 0.096470 | -1.64580184e-04 | 8.01341277e-02 | -1.76908670e+01 | 0.70 | 0 | 0 | 0 | 0 |
| M | P | 0.039564 | 0.078677 | -9.14334357e-05 | 8.01341277e-02 | -1.76908670e+01 | 0.70 | 0 | 0 | 0 | 0 |
| M | A | 0.039564 | 0.049480 | -9.14334357e-05 | 8.01341277e-02 | -1.76908670e+01 | 0.70 | -6.84807201e-05 | 5.42590026e-02 | -8.46961697e+00 | 0.70 |
| M | V | 0.039564 | 0.005578 | -9.14334357e-05 | 8.01341277e-02 | -1.76908670e+01 | 0.70 | -1.49763237e-04 | 1.15406051e-01 | -1.92322375e+01 | 0.70 |
| M | I | 0.039564 | 0.000395 | -9.14334357e-05 | 8.01341277e-02 | -1.76908670e+01 | 0.70 | -9.88175293e-05 | 1.10797972e-01 | -1.85218569e+01 | 0.70 |
| M | L | 0.039564 | 0.010998 | -9.14334357e-05 | 8.01341277e-02 | -1.76908670e+01 | 0.70 | -1.06970639e-04 | 1.17281979e-01 | -1.91871215e+01 | 0.70 |
| M | M | 0.039564 | 0.039564 | -9.14334357e-05 | 8.01341277e-02 | -1.76908670e+01 | 0.70 | -9.14334357e-05 | 8.01341277e-02 | -1.76908670e+01 | 0.70 |
| M | F | 0.039564 | 0.391642 | -9.14334357e-05 | 8.01341277e-02 | -1.76908670e+01 | 0.70 | 0 | 0 | 0 | 0 |
| M | Y | 0.039564 | 0.419186 | -9.14334357e-05 | 8.01341277e-02 | -1.76908670e+01 | 0.70 | 0 | 0 | 0 | 0 |
| M | W | 0.039564 | 0.550297 | -9.14334357e-05 | 8.01341277e-02 | -1.76908670e+01 | 0.70 | 0 | 0 | 0 | 0 |

TABLE S12: Fitness function parameters for $\varepsilon_{ij}$ where $i$ = M in Mpipi-T Model 3

| Amino Acid $i$ | Amino Acid $j$ | $\varepsilon_{ii,\text{Mpipi}}$ | $\varepsilon_{jj,\text{Mpipi}}$ | $a_i$ | $b_i$ | $c_i$ | $\alpha_i$ | $a_j$ | $b_j$ | $c_j$ | $\alpha_j$ |
|---|---|---|---|---|---|---|---|---|---|---|---|
| N | R | 0.193849 | 0.089916 | 0 | 0 | 0 | 0 | 0 | 0 | 0 | 0 |
| N | H | 0.193849 | 0.027216 | 0 | 0 | 0 | 0 | 0 | 0 | 0 | 0 |
| N | K | 0.193849 | 0.019117 | 0 | 0 | 0 | 0 | 0 | 0 | 0 | 0 |
| N | D | 0.193849 | 0.079096 | 0 | 0 | 0 | 0 | 0 | 0 | 0 | 0 |
| N | E | 0.193849 | 0.085622 | 0 | 0 | 0 | 0 | 0 | 0 | 0 | 0 |
| N | S | 0.193849 | 0.061600 | 0 | 0 | 0 | 0 | 0 | 0 | 0 | 0 |
| N | T | 0.193849 | 0.030758 | 0 | 0 | 0 | 0 | 0 | 0 | 0 | 0 |
| N | N | 0.193849 | 0.193849 | 0 | 0 | 0 | 0 | 0 | 0 | 0 | 0 |
| N | Q | 0.193849 | 0.200448 | 0 | 0 | 0 | 0 | 0 | 0 | 0 | 0 |
| N | C | 0.193849 | 0.069311 | 0 | 0 | 0 | 0 | 0 | 0 | 0 | 0 |
| N | G | 0.193849 | 0.096470 | 0 | 0 | 0 | 0 | 0 | 0 | 0 | 0 |
| N | P | 0.193849 | 0.078677 | 0 | 0 | 0 | 0 | 0 | 0 | 0 | 0 |
| N | A | 0.193849 | 0.049480 | 0 | 0 | 0 | 0 | -6.84807201e-05 | 5.42590026e-02 | -8.46961697e+00 | 0.70 |
| N | V | 0.193849 | 0.005578 | 0 | 0 | 0 | 0 | -1.49763237e-04 | 1.15406051e-01 | -1.92322375e+01 | 0.70 |
| N | I | 0.193849 | 0.000395 | 0 | 0 | 0 | 0 | -9.88175293e-05 | 1.10797972e-01 | -1.85218569e+01 | 0.70 |
| N | L | 0.193849 | 0.010998 | 0 | 0 | 0 | 0 | -1.06970639e-04 | 1.17281979e-01 | -1.91871215e+01 | 0.70 |
| N | M | 0.193849 | 0.039564 | 0 | 0 | 0 | 0 | -9.14334357e-05 | 8.01341277e-02 | -1.76908670e+01 | 0.70 |
| N | F | 0.193849 | 0.391642 | 0 | 0 | 0 | 0 | 0 | 0 | 0 | 0 |
| N | Y | 0.193849 | 0.419186 | 0 | 0 | 0 | 0 | 0 | 0 | 0 | 0 |
| N | W | 0.193849 | 0.550297 | 0 | 0 | 0 | 0 | 0 | 0 | 0 | 0 |

TABLE S13: Fitness function parameters for $\varepsilon_{ij}$ where $i$ = N in Mpipi-T Model 3

| Amino Acid $i$ | Amino Acid $j$ | $\varepsilon_{ii,\mathrm{Mpipi}}$ | $\varepsilon_{jj,\mathrm{Mpipi}}$ | $a_i$ | $b_i$ | $c_i$ | $\alpha_i$ | $a_j$ | $b_j$ | $c_j$ | $\alpha_j$ |
|---|---|---|---|---|---|---|---|---|---|---|---|
| P | R | 0.078677 | 0.089916 | 0 | 0 | 0 | 0 | 0 | 0 | 0 | 0 |
| P | H | 0.078677 | 0.027216 | 0 | 0 | 0 | 0 | 0 | 0 | 0 | 0 |
| P | K | 0.078677 | 0.019117 | 0 | 0 | 0 | 0 | 0 | 0 | 0 | 0 |
| P | D | 0.078677 | 0.079096 | 0 | 0 | 0 | 0 | 0 | 0 | 0 | 0 |
| P | E | 0.078677 | 0.085622 | 0 | 0 | 0 | 0 | 0 | 0 | 0 | 0 |
| P | S | 0.078677 | 0.061600 | 0 | 0 | 0 | 0 | 0 | 0 | 0 | 0 |
| P | T | 0.078677 | 0.030758 | 0 | 0 | 0 | 0 | 0 | 0 | 0 | 0 |
| P | N | 0.078677 | 0.193849 | 0 | 0 | 0 | 0 | 0 | 0 | 0 | 0 |
| P | Q | 0.078677 | 0.200448 | 0 | 0 | 0 | 0 | 0 | 0 | 0 | 0 |
| P | C | 0.078677 | 0.069311 | 0 | 0 | 0 | 0 | 0 | 0 | 0 | 0 |
| P | G | 0.078677 | 0.096470 | 0 | 0 | 0 | 0 | 0 | 0 | 0 | 0 |
| P | P | 0.078677 | 0.078677 | 0 | 0 | 0 | 0 | 0 | 0 | 0 | 0 |
| P | A | 0.078677 | 0.049480 | 0 | 0 | 0 | 0 | -6.84807201e-05 | 5.42590026e-02 | -8.46961697e+00 | 0.70 |
| P | V | 0.078677 | 0.005578 | 0 | 0 | 0 | 0 | -1.49763237e-04 | 1.15406051e-01 | -1.92322375e+01 | 0.70 |
| P | I | 0.078677 | 0.000395 | 0 | 0 | 0 | 0 | -9.88175293e-05 | 1.10797972e-01 | -1.85218569e+01 | 0.70 |
| P | L | 0.078677 | 0.010998 | 0 | 0 | 0 | 0 | -1.06970639e-04 | 1.17281979e-01 | -1.91871215e+01 | 0.70 |
| P | M | 0.078677 | 0.039564 | 0 | 0 | 0 | 0 | -9.14334357e-05 | 8.01341277e-02 | -1.76908670e+01 | 0.70 |
| P | F | 0.078677 | 0.391642 | 0 | 0 | 0 | 0 | 0 | 0 | 0 | 0 |
| P | Y | 0.078677 | 0.419186 | 0 | 0 | 0 | 0 | 0 | 0 | 0 | 0 |
| P | W | 0.078677 | 0.550297 | 0 | 0 | 0 | 0 | 0 | 0 | 0 | 0 |

TABLE S14: Fitness function parameters for $\varepsilon_{ij}$ where $i$ = P in Mpipi-T Model 3

| Amino Acid $i$ | Amino Acid $j$ | $\varepsilon_{ii,\mathrm{Mpipi}}$ | $\varepsilon_{jj,\mathrm{Mpipi}}$ | $a_i$ | $b_i$ | $c_i$ | $\alpha_i$ | $a_j$ | $b_j$ | $c_j$ | $\alpha_j$ |
|---|---|---|---|---|---|---|---|---|---|---|---|
| Q | R | 0.200448 | 0.089916 | 0 | 0 | 0 | 0 | 0 | 0 | 0 | 0 |
| Q | H | 0.200448 | 0.027216 | 0 | 0 | 0 | 0 | 0 | 0 | 0 | 0 |
| Q | K | 0.200448 | 0.019117 | 0 | 0 | 0 | 0 | 0 | 0 | 0 | 0 |
| Q | D | 0.200448 | 0.079096 | 0 | 0 | 0 | 0 | 0 | 0 | 0 | 0 |
| Q | E | 0.200448 | 0.085622 | 0 | 0 | 0 | 0 | 0 | 0 | 0 | 0 |
| Q | S | 0.200448 | 0.061600 | 0 | 0 | 0 | 0 | 0 | 0 | 0 | 0 |
| Q | T | 0.200448 | 0.030758 | 0 | 0 | 0 | 0 | 0 | 0 | 0 | 0 |
| Q | N | 0.200448 | 0.193849 | 0 | 0 | 0 | 0 | 0 | 0 | 0 | 0 |
| Q | Q | 0.200448 | 0.200448 | 0 | 0 | 0 | 0 | 0 | 0 | 0 | 0 |
| Q | C | 0.200448 | 0.069311 | 0 | 0 | 0 | 0 | 0 | 0 | 0 | 0 |
| Q | G | 0.200448 | 0.096470 | 0 | 0 | 0 | 0 | 0 | 0 | 0 | 0 |
| Q | P | 0.200448 | 0.078677 | 0 | 0 | 0 | 0 | 0 | 0 | 0 | 0 |
| Q | A | 0.200448 | 0.049480 | 0 | 0 | 0 | 0 | -6.84807201e-05 | 5.42590026e-02 | -8.46961697e+00 | 0.70 |
| Q | V | 0.200448 | 0.005578 | 0 | 0 | 0 | 0 | -1.49763237e-04 | 1.15406051e-01 | -1.92322375e+01 | 0.70 |
| Q | I | 0.200448 | 0.000395 | 0 | 0 | 0 | 0 | -9.88175293e-05 | 1.10797972e-01 | -1.85218569e+01 | 0.70 |
| Q | L | 0.200448 | 0.010998 | 0 | 0 | 0 | 0 | -1.06970639e-04 | 1.17281979e-01 | -1.91871215e+01 | 0.70 |
| Q | M | 0.200448 | 0.039564 | 0 | 0 | 0 | 0 | -9.14334357e-05 | 8.01341277e-02 | -1.76908670e+01 | 0.70 |
| Q | F | 0.200448 | 0.391642 | 0 | 0 | 0 | 0 | 0 | 0 | 0 | 0 |
| Q | Y | 0.200448 | 0.419186 | 0 | 0 | 0 | 0 | 0 | 0 | 0 | 0 |
| Q | W | 0.200448 | 0.550297 | 0 | 0 | 0 | 0 | 0 | 0 | 0 | 0 |

TABLE S15: Fitness function parameters for $\varepsilon_{ij}$ where $i$ = Q in Mpipi-T Model 3

| Amino Acid $i$ | Amino Acid $j$ | $\varepsilon_{ii,\text{Mpipi}}$ | $\varepsilon_{jj,\text{Mpipi}}$ | $a_i$ | $b_i$ | $c_i$ | $\alpha_i$ | $a_j$ | $b_j$ | $c_j$ | $\alpha_j$ |
|---|---|---|---|---|---|---|---|---|---|---|---|
| R | R | 0.089916 | 0.089916 | 0 | 0 | 0 | 0 | 0 | 0 | 0 | 0 |
| R | H | 0.089916 | 0.027216 | 0 | 0 | 0 | 0 | 0 | 0 | 0 | 0 |
| R | K | 0.089916 | 0.019117 | 0 | 0 | 0 | 0 | 0 | 0 | 0 | 0 |
| R | D | 0.089916 | 0.079096 | 0 | 0 | 0 | 0 | 0 | 0 | 0 | 0 |
| R | E | 0.089916 | 0.085622 | 0 | 0 | 0 | 0 | 0 | 0 | 0 | 0 |
| R | S | 0.089916 | 0.061600 | 0 | 0 | 0 | 0 | 0 | 0 | 0 | 0 |
| R | T | 0.089916 | 0.030758 | 0 | 0 | 0 | 0 | 0 | 0 | 0 | 0 |
| R | N | 0.089916 | 0.193849 | 0 | 0 | 0 | 0 | 0 | 0 | 0 | 0 |
| R | Q | 0.089916 | 0.200448 | 0 | 0 | 0 | 0 | 0 | 0 | 0 | 0 |
| R | C | 0.089916 | 0.069311 | 0 | 0 | 0 | 0 | 0 | 0 | 0 | 0 |
| R | G | 0.089916 | 0.096470 | 0 | 0 | 0 | 0 | 0 | 0 | 0 | 0 |
| R | P | 0.089916 | 0.078677 | 0 | 0 | 0 | 0 | 0 | 0 | 0 | 0 |
| R | A | 0.089916 | 0.049480 | 0 | 0 | 0 | 0 | -6.84807201e-05 | 5.42590026e-02 | -8.46961697e+00 | 0.70 |
| R | V | 0.089916 | 0.005578 | 0 | 0 | 0 | 0 | -1.49763237e-04 | 1.15406051e-01 | -1.92322375e+01 | 0.70 |
| R | I | 0.089916 | 0.000395 | 0 | 0 | 0 | 0 | -9.88175293e-05 | 1.10797972e-01 | -1.85218569e+01 | 0.70 |
| R | L | 0.089916 | 0.010998 | 0 | 0 | 0 | 0 | -1.06970639e-04 | 1.17281979e-01 | -1.91871215e+01 | 0.70 |
| R | M | 0.089916 | 0.039564 | 0 | 0 | 0 | 0 | -9.14334357e-05 | 8.01341277e-02 | -1.76908670e+01 | 0.70 |
| R | F | 0.089916 | 0.391642 | 0 | 0 | 0 | 0 | 0 | 0 | 0 | 0 |
| R | Y | 0.089916 | 0.419186 | 0 | 0 | 0 | 0 | 0 | 0 | 0 | 0 |
| R | W | 0.089916 | 0.550297 | 0 | 0 | 0 | 0 | 0 | 0 | 0 | 0 |

TABLE S16: Fitness function parameters for $\varepsilon_{ij}$ where $i$ = R in Mpipi-T Model 3

| Amino Acid $i$ | Amino Acid $j$ | $\varepsilon_{ii,\text{Mpipi}}$ | $\varepsilon_{jj,\text{Mpipi}}$ | $a_i$ | $b_i$ | $c_i$ | $\alpha_i$ | $a_j$ | $b_j$ | $c_j$ | $\alpha_j$ |
|---|---|---|---|---|---|---|---|---|---|---|---|
| S | R | 0.061600 | 0.089916 | 0 | 0 | 0 | 0 | 0 | 0 | 0 | 0 |
| S | H | 0.061600 | 0.027216 | 0 | 0 | 0 | 0 | 0 | 0 | 0 | 0 |
| S | K | 0.061600 | 0.019117 | 0 | 0 | 0 | 0 | 0 | 0 | 0 | 0 |
| S | D | 0.061600 | 0.079096 | 0 | 0 | 0 | 0 | 0 | 0 | 0 | 0 |
| S | E | 0.061600 | 0.085622 | 0 | 0 | 0 | 0 | 0 | 0 | 0 | 0 |
| S | S | 0.061600 | 0.061600 | 0 | 0 | 0 | 0 | 0 | 0 | 0 | 0 |
| S | T | 0.061600 | 0.030758 | 0 | 0 | 0 | 0 | 0 | 0 | 0 | 0 |
| S | N | 0.061600 | 0.193849 | 0 | 0 | 0 | 0 | 0 | 0 | 0 | 0 |
| S | Q | 0.061600 | 0.200448 | 0 | 0 | 0 | 0 | 0 | 0 | 0 | 0 |
| S | C | 0.061600 | 0.069311 | 0 | 0 | 0 | 0 | 0 | 0 | 0 | 0 |
| S | G | 0.061600 | 0.096470 | 0 | 0 | 0 | 0 | 0 | 0 | 0 | 0 |
| S | P | 0.061600 | 0.078677 | 0 | 0 | 0 | 0 | 0 | 0 | 0 | 0 |
| S | A | 0.061600 | 0.049480 | 0 | 0 | 0 | 0 | -6.84807201e-05 | 5.42590026e-02 | -8.46961697e+00 | 0.70 |
| S | V | 0.061600 | 0.005578 | 0 | 0 | 0 | 0 | -1.49763237e-04 | 1.15406051e-01 | -1.92322375e+01 | 0.70 |
| S | I | 0.061600 | 0.000395 | 0 | 0 | 0 | 0 | -9.88175293e-05 | 1.10797972e-01 | -1.85218569e+01 | 0.70 |
| S | L | 0.061600 | 0.010998 | 0 | 0 | 0 | 0 | -1.06970639e-04 | 1.17281979e-01 | -1.91871215e+01 | 0.70 |
| S | M | 0.061600 | 0.039564 | 0 | 0 | 0 | 0 | -9.14334357e-05 | 8.01341277e-02 | -1.76908670e+01 | 0.70 |
| S | F | 0.061600 | 0.391642 | 0 | 0 | 0 | 0 | 0 | 0 | 0 | 0 |
| S | Y | 0.061600 | 0.419186 | 0 | 0 | 0 | 0 | 0 | 0 | 0 | 0 |
| S | W | 0.061600 | 0.550297 | 0 | 0 | 0 | 0 | 0 | 0 | 0 | 0 |

TABLE S17: Fitness function parameters for $\varepsilon_{ij}$ where $i$ = S in Mpipi-T Model 3

| Amino Acid $i$ | Amino Acid $j$ | $\varepsilon_{ii,\text{Mpipi}}$ | $\varepsilon_{jj,\text{Mpipi}}$ | $a_i$ | $b_i$ | $c_i$ | $\alpha_i$ | $a_j$ | $b_j$ | $c_j$ | $\alpha_j$ |
|---|---|---|---|---|---|---|---|---|---|---|---|
| T | R | 0.030758 | 0.089916 | 0 | 0 | 0 | 0 | 0 | 0 | 0 | 0 |
| T | H | 0.030758 | 0.027216 | 0 | 0 | 0 | 0 | 0 | 0 | 0 | 0 |
| T | K | 0.030758 | 0.019117 | 0 | 0 | 0 | 0 | 0 | 0 | 0 | 0 |
| T | D | 0.030758 | 0.079096 | 0 | 0 | 0 | 0 | 0 | 0 | 0 | 0 |
| T | E | 0.030758 | 0.085622 | 0 | 0 | 0 | 0 | 0 | 0 | 0 | 0 |
| T | S | 0.030758 | 0.061600 | 0 | 0 | 0 | 0 | 0 | 0 | 0 | 0 |
| T | T | 0.030758 | 0.030758 | 0 | 0 | 0 | 0 | 0 | 0 | 0 | 0 |
| T | N | 0.030758 | 0.193849 | 0 | 0 | 0 | 0 | 0 | 0 | 0 | 0 |
| T | Q | 0.030758 | 0.200448 | 0 | 0 | 0 | 0 | 0 | 0 | 0 | 0 |
| T | C | 0.030758 | 0.069311 | 0 | 0 | 0 | 0 | 0 | 0 | 0 | 0 |
| T | G | 0.030758 | 0.096470 | 0 | 0 | 0 | 0 | 0 | 0 | 0 | 0 |
| T | P | 0.030758 | 0.078677 | 0 | 0 | 0 | 0 | 0 | 0 | 0 | 0 |
| T | A | 0.030758 | 0.049480 | 0 | 0 | 0 | 0 | -6.84807201e-05 | 5.42590026e-02 | -8.46961697e+00 | 0.70 |
| T | V | 0.030758 | 0.005578 | 0 | 0 | 0 | 0 | -1.49763237e-04 | 1.15406051e-01 | -1.92322375e+01 | 0.70 |
| T | I | 0.030758 | 0.000395 | 0 | 0 | 0 | 0 | -9.88175293e-05 | 1.10797972e-01 | -1.85218569e+01 | 0.70 |
| T | L | 0.030758 | 0.010998 | 0 | 0 | 0 | 0 | -1.06970639e-04 | 1.17281979e-01 | -1.91871215e+01 | 0.70 |
| T | M | 0.030758 | 0.039564 | 0 | 0 | 0 | 0 | -9.14334357e-05 | 8.01341277e-02 | -1.76908670e+01 | 0.70 |
| T | F | 0.030758 | 0.391642 | 0 | 0 | 0 | 0 | 0 | 0 | 0 | 0 |
| T | Y | 0.030758 | 0.419186 | 0 | 0 | 0 | 0 | 0 | 0 | 0 | 0 |
| T | W | 0.030758 | 0.550297 | 0 | 0 | 0 | 0 | 0 | 0 | 0 | 0 |

TABLE S18: Fitness function parameters for $\varepsilon_{ij}$ where $i$ = T in Mpipi-T Model 3

| Amino Acid $i$ | Amino Acid $j$ | $\varepsilon_{ii,\text{Mpipi}}$ | $\varepsilon_{jj,\text{Mpipi}}$ | $a_i$ | $b_i$ | $c_i$ | $\alpha_i$ | $a_j$ | $b_j$ | $c_j$ | $\alpha_j$ |
|---|---|---|---|---|---|---|---|---|---|---|---|
| V | R | 0.005578 | 0.089916 | -1.49763237e-04 | 1.15406051e-01 | -1.92322375e+01 | 0.70 | 0 | 0 | 0 | 0 |
| V | H | 0.005578 | 0.027216 | -1.49763237e-04 | 1.15406051e-01 | -1.92322375e+01 | 0.70 | 0 | 0 | 0 | 0 |
| V | K | 0.005578 | 0.019117 | -1.49763237e-04 | 1.15406051e-01 | -1.92322375e+01 | 0.70 | 0 | 0 | 0 | 0 |
| V | D | 0.005578 | 0.079096 | -1.49763237e-04 | 1.15406051e-01 | -1.92322375e+01 | 0.70 | 0 | 0 | 0 | 0 |
| V | E | 0.005578 | 0.085622 | -1.49763237e-04 | 1.15406051e-01 | -1.92322375e+01 | 0.70 | 0 | 0 | 0 | 0 |
| V | S | 0.005578 | 0.061600 | -1.49763237e-04 | 1.15406051e-01 | -1.92322375e+01 | 0.70 | 0 | 0 | 0 | 0 |
| V | T | 0.005578 | 0.030758 | -1.49763237e-04 | 1.15406051e-01 | -1.92322375e+01 | 0.70 | 0 | 0 | 0 | 0 |
| V | N | 0.005578 | 0.193849 | -1.49763237e-04 | 1.15406051e-01 | -1.92322375e+01 | 0.70 | 0 | 0 | 0 | 0 |
| V | Q | 0.005578 | 0.200448 | -1.49763237e-04 | 1.15406051e-01 | -1.92322375e+01 | 0.70 | 0 | 0 | 0 | 0 |
| V | C | 0.005578 | 0.069311 | -1.49763237e-04 | 1.15406051e-01 | -1.92322375e+01 | 0.70 | 0 | 0 | 0 | 0 |
| V | G | 0.005578 | 0.096470 | -2.69573827e-04 | 1.15406051e-01 | -1.92322375e+01 | 0.70 | 0 | 0 | 0 | 0 |
| V | P | 0.005578 | 0.078677 | -1.49763237e-04 | 1.15406051e-01 | -1.92322375e+01 | 0.70 | 0 | 0 | 0 | 0 |
| V | A | 0.005578 | 0.049480 | -1.49763237e-04 | 1.15406051e-01 | -1.92322375e+01 | 0.70 | -6.84807201e-05 | 5.42590026e-02 | -8.46961697e+00 | 0.70 |
| V | V | 0.005578 | 0.005578 | -1.49763237e-04 | 1.15406051e-01 | -1.92322375e+01 | 0.70 | -1.49763237e-04 | 1.15406051e-01 | -1.92322375e+01 | 0.70 |
| V | I | 0.005578 | 0.000395 | -1.49763237e-04 | 1.15406051e-01 | -1.92322375e+01 | 0.70 | -9.88175293e-05 | 1.10797972e-01 | -1.85218569e+01 | 0.70 |
| V | L | 0.005578 | 0.010998 | -1.49763237e-04 | 1.15406051e-01 | -1.92322375e+01 | 0.70 | -1.06970639e-04 | 1.17281979e-01 | -1.91871215e+01 | 0.70 |
| V | M | 0.005578 | 0.039564 | -1.49763237e-04 | 1.15406051e-01 | -1.92322375e+01 | 0.70 | -9.14334357e-05 | 8.01341277e-02 | -1.76908670e+01 | 0.70 |
| V | F | 0.005578 | 0.391642 | -1.49763237e-04 | 1.15406051e-01 | -1.92322375e+01 | 0.70 | 0 | 0 | 0 | 0 |
| V | Y | 0.005578 | 0.419186 | -1.49763237e-04 | 1.15406051e-01 | -1.92322375e+01 | 0.70 | 0 | 0 | 0 | 0 |
| V | W | 0.005578 | 0.550297 | -1.49763237e-04 | 1.15406051e-01 | -1.92322375e+01 | 0.70 | 0 | 0 | 0 | 0 |

TABLE S19: Fitness function parameters for $\varepsilon_{ij}$ where $i$ = V in Mpipi-T Model 3

| Amino Acid $i$ | Amino Acid $j$ | $\varepsilon_{ii,\mathrm{Mpipi}}$ | $\varepsilon_{jj,\mathrm{Mpipi}}$ | $a_i$ | $b_i$ | $c_i$ | $\alpha_i$ | $a_j$ | $b_j$ | $c_j$ | $\alpha_j$ |
|---|---|---|---|---|---|---|---|---|---|---|---|
| W | R | 0.550297 | 0.089916 | 0 | 0 | 0 | 0 | 0 | 0 | 0 | 0 |
| W | H | 0.550297 | 0.027216 | 0 | 0 | 0 | 0 | 0 | 0 | 0 | 0 |
| W | K | 0.550297 | 0.019117 | 0 | 0 | 0 | 0 | 0 | 0 | 0 | 0 |
| W | D | 0.550297 | 0.079096 | 0 | 0 | 0 | 0 | 0 | 0 | 0 | 0 |
| W | E | 0.550297 | 0.085622 | 0 | 0 | 0 | 0 | 0 | 0 | 0 | 0 |
| W | S | 0.550297 | 0.061600 | 0 | 0 | 0 | 0 | 0 | 0 | 0 | 0 |
| W | T | 0.550297 | 0.030758 | 0 | 0 | 0 | 0 | 0 | 0 | 0 | 0 |
| W | N | 0.550297 | 0.193849 | 0 | 0 | 0 | 0 | 0 | 0 | 0 | 0 |
| W | Q | 0.550297 | 0.200448 | 0 | 0 | 0 | 0 | 0 | 0 | 0 | 0 |
| W | C | 0.550297 | 0.069311 | 0 | 0 | 0 | 0 | 0 | 0 | 0 | 0 |
| W | G | 0.550297 | 0.096470 | 0 | 0 | 0 | 0 | 0 | 0 | 0 | 0 |
| W | P | 0.550297 | 0.078677 | 0 | 0 | 0 | 0 | 0 | 0 | 0 | 0 |
| W | A | 0.550297 | 0.049480 | 0 | 0 | 0 | 0 | -6.84807201e-05 | 5.42590026e-02 | -8.46961697e+00 | 0.70 |
| W | V | 0.550297 | 0.005578 | 0 | 0 | 0 | 0 | -1.49763237e-04 | 1.15406051e-01 | -1.92322375e+01 | 0.70 |
| W | I | 0.550297 | 0.000395 | 0 | 0 | 0 | 0 | -9.88175293e-05 | 1.10797972e-01 | -1.85218569e+01 | 0.70 |
| W | L | 0.550297 | 0.010998 | 0 | 0 | 0 | 0 | -1.06970639e-04 | 1.17281979e-01 | -1.91871215e+01 | 0.70 |
| W | M | 0.550297 | 0.039564 | 0 | 0 | 0 | 0 | -9.14334357e-05 | 8.01341277e-02 | -1.76908670e+01 | 0.70 |
| W | F | 0.550297 | 0.391642 | 0 | 0 | 0 | 0 | 0 | 0 | 0 | 0 |
| W | Y | 0.550297 | 0.419186 | 0 | 0 | 0 | 0 | 0 | 0 | 0 | 0 |
| W | W | 0.550297 | 0.550297 | 0 | 0 | 0 | 0 | 0 | 0 | 0 | 0 |

TABLE S20: Fitness function parameters for $\varepsilon_{ij}$ where $i$ = W in Mpipi-T Model 3

| Amino Acid $i$ | Amino Acid $j$ | $\varepsilon_{ii,\mathrm{Mpipi}}$ | $\varepsilon_{jj,\mathrm{Mpipi}}$ | $a_i$ | $b_i$ | $c_i$ | $\alpha_i$ | $a_j$ | $b_j$ | $c_j$ | $\alpha_j$ |
|---|---|---|---|---|---|---|---|---|---|---|---|
| Y | R | 0.419186 | 0.089916 | 0 | 0 | 0 | 0 | 0 | 0 | 0 | 0 |
| Y | H | 0.419186 | 0.027216 | 0 | 0 | 0 | 0 | 0 | 0 | 0 | 0 |
| Y | K | 0.419186 | 0.019117 | 0 | 0 | 0 | 0 | 0 | 0 | 0 | 0 |
| Y | D | 0.419186 | 0.079096 | 0 | 0 | 0 | 0 | 0 | 0 | 0 | 0 |
| Y | E | 0.419186 | 0.085622 | 0 | 0 | 0 | 0 | 0 | 0 | 0 | 0 |
| Y | S | 0.419186 | 0.061600 | 0 | 0 | 0 | 0 | 0 | 0 | 0 | 0 |
| Y | T | 0.419186 | 0.030758 | 0 | 0 | 0 | 0 | 0 | 0 | 0 | 0 |
| Y | N | 0.419186 | 0.193849 | 0 | 0 | 0 | 0 | 0 | 0 | 0 | 0 |
| Y | Q | 0.419186 | 0.200448 | 0 | 0 | 0 | 0 | 0 | 0 | 0 | 0 |
| Y | C | 0.419186 | 0.069311 | 0 | 0 | 0 | 0 | 0 | 0 | 0 | 0 |
| Y | G | 0.419186 | 0.096470 | 0 | 0 | 0 | 0 | 0 | 0 | 0 | 0 |
| Y | P | 0.419186 | 0.078677 | 0 | 0 | 0 | 0 | 0 | 0 | 0 | 0 |
| Y | A | 0.419186 | 0.049480 | 0 | 0 | 0 | 0 | -6.84807201e-05 | 5.42590026e-02 | -8.46961697e+00 | 0.70 |
| Y | V | 0.419186 | 0.005578 | 0 | 0 | 0 | 0 | -1.49763237e-04 | 1.15406051e-01 | -1.92322375e+01 | 0.70 |
| Y | I | 0.419186 | 0.000395 | 0 | 0 | 0 | 0 | -9.88175293e-05 | 1.10797972e-01 | -1.85218569e+01 | 0.70 |
| Y | L | 0.419186 | 0.010998 | 0 | 0 | 0 | 0 | -1.06970639e-04 | 1.17281979e-01 | -1.91871215e+01 | 0.70 |
| Y | M | 0.419186 | 0.039564 | 0 | 0 | 0 | 0 | -9.14334357e-05 | 8.01341277e-02 | -1.76908670e+01 | 0.70 |
| Y | F | 0.419186 | 0.391642 | 0 | 0 | 0 | 0 | 0 | 0 | 0 | 0 |
| Y | Y | 0.419186 | 0.419186 | 0 | 0 | 0 | 0 | 0 | 0 | 0 | 0 |
| Y | W | 0.419186 | 0.550297 | 0 | 0 | 0 | 0 | 0 | 0 | 0 | 0 |

TABLE S21: Fitness function parameters for $\varepsilon_{ij}$ where $i$ = Y in Mpipi-T Model 3

[1] F. G. Quiroz and A. Chilkoti, Sequence heuristics to encode phase behaviour in intrinsically disordered protein polymers, Nature Materials **14**, 1164 (2015).
[2] J. A. Joseph, A. Reinhardt, A. Aguirre, P. Y. Chew, K. O. Russell, J. R. Espinosa, A. Garaizar, and R. Collepardo-Guevara, Physics-driven coarse-grained model for biomolecular phase separation with near-quantitative accuracy, Nature Computational Science **1**, 732 (2021).
[3] A. Bremer, M. Farag, W. M. Borcherds, I. Peran, E. W. Martin, R. V. Pappu, and T. Mittag, Deciphering how naturally occurring sequence features impact the phase behaviours of disordered prion-like domains, Nature Chemistry **14**, 196 (2022).
[4] J. R. Simon, N. J. Carroll, M. Rubinstein, A. Chilkoti, and G. P. López, Programming molecular self-assembly of intrinsically disordered proteins containing sequences of low complexity, Nature Chemistry **9**, 509 (2017).
